# Multiple scenarios for sexual crosses in the fungal pathogen *Zymoseptoria tritici* on wheat residues: potential consequences for virulence gene transmission

**DOI:** 10.1101/2022.02.24.481803

**Authors:** Carolina Orellana-Torrejon, Tiphaine Vidal, Gwilherm Gazeau, Anne-Lise Boixel, Sandrine Gélisse, Jérôme Lageyre, Sébastien Saint-Jean, Frédéric Suffert

**Affiliations:** Université Paris-Saclay, INRAE, UR BIOGER, 78850 Thiverval-Grignon, France; Université Paris-Saclay, INRAE, AgroParisTech, UMR ECOSYS, 78850 Thiverval-Grignon, France

**Keywords:** asexual multiplication, avirulence, infection, plant disease epidemiology, sexual reproduction, resistance breakdown

## Abstract

Little is known about the impact of host immunity on sexual reproduction in fungal pathogens. In particular, it is unclear whether crossing requires both sexual partners to infect living plant tissues. We addressed this issue in a three-year experiment investigating different scenarios of *Zymoseptoria tritici* crosses on wheat according to the virulence (‘vir’) or avirulence (‘avr’) of the parents against a qualitative resistance gene. Co-inoculations (‘vir × vir’, ‘avr × vir’, ‘avr × avr’) and single inoculations were performed on a cultivar carrying the resistance gene (Cellule) and a susceptible cultivar (Apache), in the greenhouse. We assessed the intensity of asexual multiplication by scoring disease severity, and the intensity of sexual reproduction by counting the ascospores discharged from wheat residues. As expected, disease severity was more intense on Cellule for ‘vir × vir’ co-inoculations than for ‘avr × vir’ co-inoculations, with no disease for ‘avr × avr’. However, all types of co-inoculation yielded sexual offspring, whether or not the parental strains caused plant symptoms. Parenthood was confirmed by genotyping (SSR markers), and the occurrence of crosses between (co-)inoculated and exogenous strains (other strains from the experiment, or from far away) was determined. We found that symptomatic asexual infection was not required for a strain to participate in sexual reproduction, and that avirulent strains could be maintained asymptomatically “on” or “in” leaf tissues of plants carrying the corresponding resistant gene for long enough to reproduce sexually. In two of the three years, the intensity of sexual reproduction did not differ significantly between the three types of co-inoculation in Cellule, suggesting that crosses involving avirulent strains are not anecdotal. We discuss the possible mechanisms explaining the maintenance of avirulence in *Z. tritici* populations and supporting the potential efficacy of cultivar mixtures for limiting resistance gene breakdown.

**Highlights:**

- Avirulent *Zymoseptoria tritici* strains can reproduce sexually in wheat plants carrying the corresponding resistant gene.
- Symptomatic infection of plant tissues is not essential for a strain to reproduce sexually.
- Avirulent strains can be maintained asymptomatically “on” or “in” leaf tissues of plants carrying the corresponding resistant gene for long enough to reproduce sexually.
- Crosses of virulent strains with virulent and avirulent strains in a plant host carrying the corresponding resistance gene can produce offspring with similar population sizes.
- Several possible scenarios for sexual crosses can explain the maintenance of avirulence in *Zymoseptoria tritici* populations evolving in a wheat canopy, particular in cultivar mixtures.

## 1. Introduction

Sexual reproduction drives genetic recombination in many phytopathogenic fungal populations (Ni *et al*., 2011). It also plays an essential role in their interseason survival and, thus, in the multiannual recurrence of many plant disease epidemics (Burdon and Laine, 2019). It is widely believed that sexual reproduction in foliar pathogens requires “infection” — penetration and growth in the organs of a living host plant, including the appearance of symptoms — by both individual partners during an epidemic. Some host plants carry qualitative resistance genes that prevent or stop infection by pathogenic strains that are “avirulent” (‘avr’), according to the “gene-for-gene” model (Dangl and Jones, 2001). Pathogenic “virulent” strains (‘vir’) able to infect these “resistant” plants can then reproduce sexually in host tissues, whereas avirulent strains, having been prevented from infecting the host plants, are presumed not to be able to engage in sexual reproduction. Avirulent strains are, therefore, thought to be progressively eliminated from host plans populations carrying the corresponding resistance gene. However, this assumption is simplistic, because the absence of infection (and, thus, of symptoms) does not necessarily prevent a physical encounter between sexual partners. Indeed, many fungal species reported to act as plant pathogens can also develop epiphytically, endophytically (causing an “asymptomatic infection”), or as saprotrophs on plant residues (Kerdraon *et al*., 2019; Selosse *et al*., 2018; Wheeler *et al*., 2019). As pathogens can complete their life cycle without necessarily causing symptoms, it may be possible for avirulent strains to reproduce sexually on a host that they failed to infect. Clarification of this point is essential, not only from the standpoint of fungal genetics and biology, but also for epidemiology, as ‘unexpected’ sexual reproduction events may affect the ratio of virulent to avirulent strains in the offspring population developing on a cultivar considered resistant. Such events could affect the efficacy of the resistance gene and the durability of resistance deployment strategies.

There are still many gaps in our knowledge of the interactions between living plants and fungi during sexual reproduction in natural conditions, because of experimental difficulties promoting sexual reproduction in some of the organisms studied, and because of their ability to switch between lifestyles — epiphytic, endophytic and pathogenic — under changing environmental conditions (e.g., Schulz and Boyle, 2005). Sexual reproduction in several fungi can be achieved under controlled conditions *in planta*, but also sometimes in dead plant tissues colonised in a saprophytic way, or in axenic conditions *in vitro*. The ease of crossing is heterogeneous in heterothallic species, which require two sexually compatible parental strains. For example, crosses were obtained for *Phaeosphaeria nodorum* on wheat residues (straw) by Halama & Lacoste (1992), but the same authors were unable to obtain crosses of *Zymoseptoria tritici*, the causal agent of Septoria tritici blotch (STB) in wheat under similar conditions. These difficulties do not necessarily reflect what happens in natural conditions. In several ascomycetes, the sexual form is cryptic or facultative, and therefore difficult to detect. For instance, the sexual form has never been identified for the rice pathogen *Magnaporthe oryzae* in natural conditions, whereas it can be obtained *in vitro* (Saleh *et al*., 2012a, 2012b). Controlled crosses of *Venturia inaequalis* were first obtained on apple trees in the greenhouse by Keitt & Palmiter (1937), then on detached apple leaf discs by Ross & Hamlin (1962) and Martin *et al*. (1981), and, later, on artificial medium by Sierotzki & Gessler (1998). The causal agent of oilseed rape stem canker, *Leptosphaeria maculans,* was successfully crossed on artificial media (Shoemaker and Brun, 2011), but sexual reproduction *in planta* under controlled conditions was not achieved until recently (Bousset *et al*., 2018). The physiological and cellular mechanisms occurring during sexual reproduction have been described in some model ascomycetes (e.g. *Neurospora crassa*; Brun *et al*., 2021), but the interaction with the host plant remains poorly understood for biotrophic and hemibiotrophic plant pathogens.

Sexual reproduction helps maintain pathogen populations between epidemics and is involved in the interseason transmission of primary inoculum. It is, thus, essential to understand how plant host immunity affects this process and its consequences for a local pathogen population. The efficacy of strategies for controlling fungal plant pathogens through cultivar resistance is generally assessed at cropping-season scale. For this reason, the impact of host “immunity” on the sexual reproduction of the pathogen, which mainly occurs between cropping seasons, has been much less studied. Host-plant resistance could be considered as a form of stress, in which case it would be expected to increase sexual reproduction rates. However, contrasting results have been obtained for pathosystems in which this question has been addressed. *V. inaequalis* has been reported to grow and produce sexual fruiting bodies (pseudothecia) on pear and apple leaves, but its growth on different apple cultivars was not related to the susceptibility of these cultivars to apple scab (Ross and Hamlin, 1962). Conversely, Marcroft *et al*. (2004), Bousset *et al*. (2021) and Fortune *et al*. (2021) highlighted a significant effect of oilseed rape genotype on the density of *L. maculans* pseudothecia produced on crop residues. The intensity of sexual reproduction was shown to depend on disease severity at harvest, with higher stem canker severities leading to the production of more pseudothecia. A similar finding was reported for the hemibiotrophic pathogen *Z. tritici*: a positive correlation was found between the susceptibility of wheat cultivars, expressed as disease severity during the epidemic stage, and the intensity of sexual reproduction on wheat residues (Cowger *et al*., 2002). Such a correlation is not surprising, given that this fungus is heterothallic, the opposite mating types must come into contact for sex to occur (Kema *et al*., 1996), and the two mating-type idiomorphs are consistently found to be evenly distributed, even at a fine scale (plant and field; Zhan *et al*., 2002; Siah *et al*., 2010; Morais *et al*., 2019). However, data are lacking as to whether there is systematically a good match between the levels of host susceptibility at the asexual and sexual stages (Suffert *et al*., 2016), apart from the empirical characterisation of density-dependent processes established by Suffert *et al*. (2018a). Moreover, it has been shown that a few days’ difference in the latent periods of the strains, with one progressing faster than the other in wheat tissues, is slightly beneficial to ascosporogenesis, whereas a two- to three-week interval between infections with the two strains is detrimental (Suffert *et al*., 2016). More generally, the conditions of asexual infection by two parental strains of *Z. tritici* affect the dynamics of sexual reproduction, but it remains challenging to identify these conditions and to decipher the mechanisms involved.

Genes conferring partial or complete resistance to an ascomycete pathogen on the host plant may also influence the intensity of the pathogen’s sexual reproduction, thereby limiting the density of sexual fruiting bodies and/or ascosporogenesis. The production and release of smaller numbers of ascospores may be another immunity trait lowering disease severity during the early stages of the subsequent epidemic at least (e.g. Suffert and Sache, 2011), and ensuring better crop yields. This is an important point consider, although not the focus of this study. It would have epidemiological consequences, as highlighted by the trade-offs identified between crop infection and survival or the interseason transmission of primary inoculum in *Phytophthora infestans* (Pasco *et al*., 2016), *Plasmopara viticola* (Delbac *et al*., 2019) and *Z. tritici* (Suffert *et al*., 2018b).

STB, one of the most important wheat diseases worldwide, is a suitable biological model for the investigation of these issues. The presence of *Stb* genes conferring resistance in wheat cultivars (Brown *et al*., 2015) is known to determine the ability of virulent and avirulent strains to infect them. If we assume that only virulent strains can cross on a cultivar carrying a resistance gene, then the offspring would be exclusively virulent, resulting in the fixation of this virulence at a very high frequency, especially when the *Stb* gene is present in most, if not all of the cultivars widely deployed in a wheat production area. However, the epidemiological situation is more complex in the case of *Z. tritici*. For instance, the frequency of strains avirulent against *Stb6* remains at about 0.1, despite the widespread use of this resistance gene in wheat cultivars deployed in France (Saintenac *et al*., 2018; Stephens *et al*., 2021). Various processes, such as the possibility of ‘avr × vir’ crosses on a cultivar carrying *Stb6,* could counter the generalisation of virulence. Indeed, Kema *et al*. (2018) established experimentally that such a cross can occur between a *Z. tritici* virulent strain successfully infecting host seedlings carrying the corresponding resistance gene and an avirulent strain unable to infect the host. This result should be interpreted in light of the ‘epiphytic’ growth of *Z. tritici*, which has recently been shown to occur for at least 10 days in specific humidity conditions (Fones *et al*., 2017). A second potential mechanism should also be considered: the facilitation between co-infecting fungal strains demonstrated by Dutt *et al*. (2021), after a ‘systemic induced susceptibility’ (SIS) reaction (Seybold *et al*., 2020) or ‘neighbour-modulated susceptibility’ (NMS) reaction (Pélissier *et al*., 2021) for instance, in *Z. tritici*, with consequences modelled by Sofonea *et al*. (2017). These mechanisms, explaining the possible simultaneous co-infection of a plant by an avirulent and a virulent strain, are consistent with the presence of avirulent strains in STB lesions sampled from resistant cultivars (e.g. Orellana-Torrejon *et al*., 2022a). Some authors have also suggested the possible induction of ‘systemic acquired resistance’ (SAR) or ‘localised acquired resistance’ (LAR) in other pathosystems (Costet *et al*., 1999), although their studies focused on levels of aggressiveness (quantitative response) rather than virulence (qualitative response). Microscopy studies showed, for example, that the *Stb16q* gene stops the progression of avirulent *Z. tritici* strains 10 day after inoculation, either before their penetration through the stomata or in the substomatal cavities (Saintenac *et al*., 2021). The physiological processes induced in such incompatible reactions merit exploration, as they may also explain the phenomenon of partial resistance against co-inoculation with virulent and avirulent strains. They could potentially decrease the severity of STB severity observed in co-infected wheat plants (Barrett *et al*., 2021; Schürch and Roy, 2004). Other mechanisms should also be considered, such as crosses between non-infecting strains developing saprophytically on (or in) wheat residues (i.e. dead tissues). Crossing in such conditions has been reported for *P. nodorum* but has never been observed for *Z. tritici* (Halama & Lacoste, 1992).

In this experimental study, we investigated the diversity of scenarios in which *Z. tritici* sexual reproduction could occur in a wheat cultivar carrying an *Stb* resistance gene. Our goal was to determine whether: (i) strains that fail to infect wheat tissues during the asexual phase can reproduce sexually (“Who?”); (ii) sex can occur if one of the parental strains arrives on plant tissues later than the other, particularly after the senescence of these tissues (“When?”); (iii) the presence of parental strains in different proportions affects the intensity of sexual reproduction (“How?”). As a means of addressing the question “Do these situations actually lead to sexual crosses?”, we designed an experiment based on different crossing situations, with three types of co-inoculation (‘vir × vir’, ‘avr × vir’ and ‘avr × avr’) and the corresponding single inoculations (virulent, avirulent), involving multiple epidemiological processes (spore transfers between different types of wheat tissues and fungal growth conditions). Obtaining an answer to this question implied finding a balance between natural conditions allowing the processes to be expressed and controlled conditions managing and dissociating the processes considered, at least partially. We, thus, simplified the host-pathogen interactions, using only pairs of parental strains and two wheat cultivars, for which virulence (vir, avr) and resistance (R, S) status, respectively, were known.

## 2. Materials & methods

### 2.1. Overall strategy

In this experiment, we inoculated adult wheat plants with and without a resistance gene with several virulent and avirulent *Z. tritici* strains in the greenhouse, to estimate the intensity of sexual reproduction by quantifying the offspring. We performed ‘co-inoculations’ with pairs of compatible parental strains and ‘single inoculations’ of each parent, to estimate the possibility of a strain crossing with an exogenous strain (used for another cross during the same experiment or from outside) in the various crossing situations. The experiment was replicated over three years (2018, 2019 and 2020). Each year, we performed 12 co-inoculations with eight parental strains, and eight single inoculations (**Figure 1**). We tested three types of cross, defined in terms of the virulence status of the parental strains: both virulent (‘vir × vir’), both avirulent (‘avr × avr’), and one virulent and the other avirulent (‘vir × avr’). All co-inoculations were performed with suspensions of equal proportions of blastospores from the two parental strains sprayed on adult wheat plants. Each year, additional co-inoculations were performed for one of the ‘avr × vir’ treatments, with unbalanced proportions for the biparental suspensions. Disease severity was assessed to estimate the intensity of asexual multiplication. Plants were left in the greenhouse and allowed to follow their natural development cycle until they had completely dried out. The resulting senescent plants, which were considered to be ‘residues’, were placed outdoors for several months, and the intensity of sexual reproduction was then estimated by quantifying ascospore discharge, as described by Suffert *et al*. (2016), for each crossing situation. Several offspring strains were sampled from some sets of residues and genotyped with 12 SSR markers (Gautier *et al*., 2014) to check their ‘parenthood’: the genotypic profiles of the offspring were compared with those of the (co-)inoculated strains to determine which strains had actually crossed, their status (virulent and/or avirulent) and their origin (inoculated and/or exogenous).

**Figure 1.**
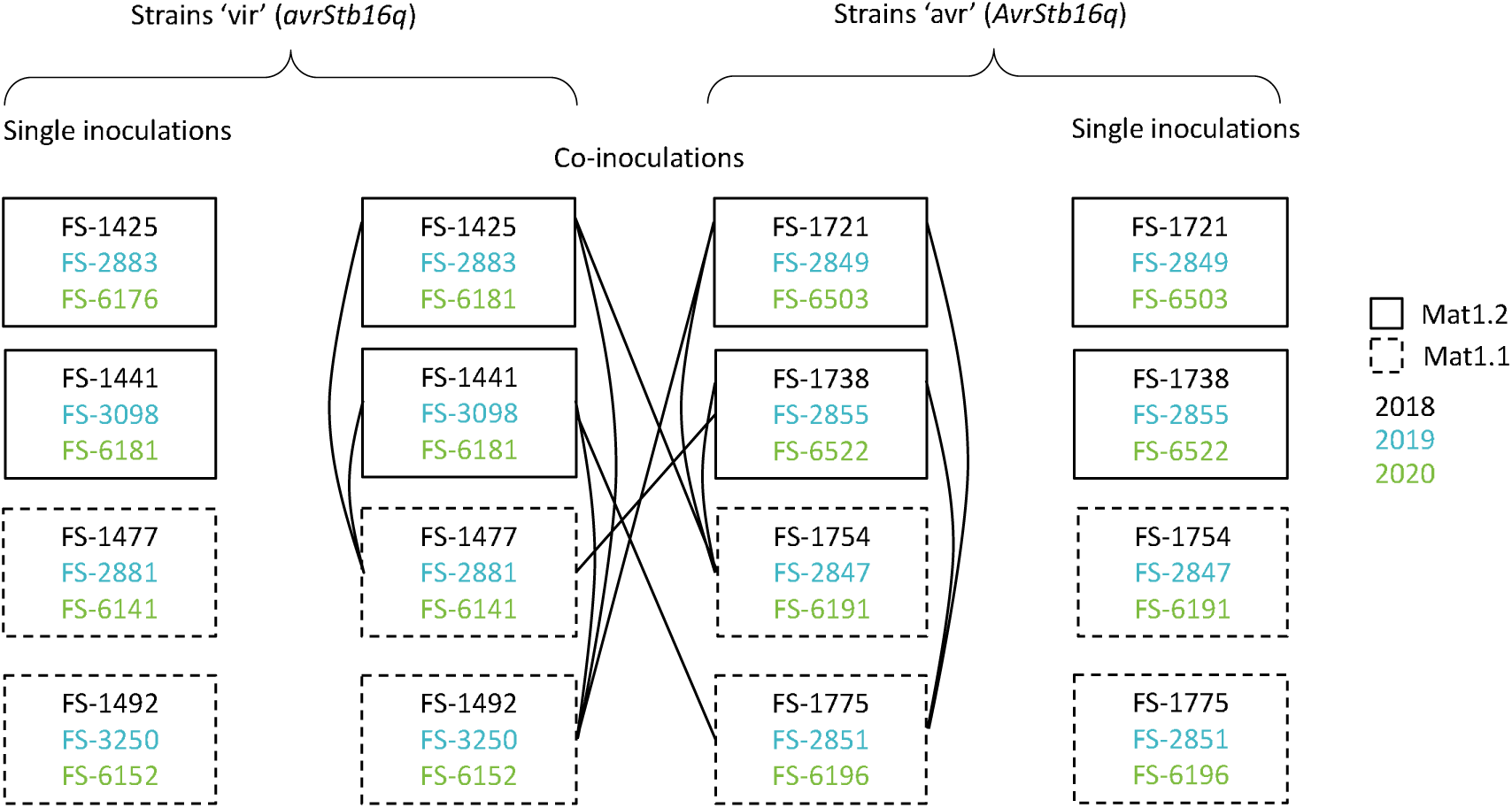
Design of the three-year experiment (2018, 2019, 2020) involving *Zymoseptoria tritici* strains virulent (‘vir’) and avirulent (‘avr’) against *Stb16q*. Each adult wheat plant of cv. Apache or Cellule received the same number of blastospores, whether inoculated with one (eight single inoculations) or two strains (12 co-inoculations). All co-inoculations were performed with equiproportional biparental inoculum suspensions. For one of the ‘avr × vir’ co-inoculations (FS1425 × FS1754 in 2018, FS2883 × FS2847 in 2019, FS6176 × FS6191 in 2020), four additional co-inoculations were performed with biparental suspensions in unbalanced proportions (0.1-0.9, 0.25-0.75, 0.75-0.25, and 0.9-0.1).

### 2.2. Plant material

We used cv. Cellule (Florimond Desprez, France) carrying the *Stb16q* resistance gene (Ghaffary *et al*., 2012) for its resistance to STB and Apache (Limagrain Europe, France) for its susceptibility. Both cultivars have been widely grown in France in the last decade. A few years after the deployment of Cellule (2012), strains capable of overcoming *Stb16q* were detected. These strains now comprise a substantial proportion of *Z. tritici* populations in Europe (Kildea *et al*., 2020; Orellana-Torrejon *et al*., 2022a). Seeds of cv. Cellule and Apache were sown on 11^th^ December 2018, 20^th^ December 2019, and 9^th^ December 2020, in peat pots (Jiffy). Seedlings were kept in greenhouse conditions for two weeks and were then vernalised in a growth chamber for 8 weeks at 8°C, with a 10-h light/14-h dark photoperiod. The plants were then returned to the greenhouse and left to acclimate for one week before transplantation into individual pots filled with 1 litre of commercial compost (Klasmann Peat Substrate 4; Klasmann France SARL, France) supplemented with 4 g of slow-release fertiliser (Osmocote Exact 16-11-11N-P-K 3MgO Te) per pot. The plants were also watered with Hydrokani C2 fertiliser (Hydro Agri Spécialités, France) diluted 1:100 and poured into the saucers under the pots. During the growth period, the plants were illuminated with natural daylight supplemented with 400 W sodium vapour lamps. The air temperature was kept below 20°C during the 15-hour light period and above 15°C during the 9-hour dark period. Plants were thinned to three stems per pot during the growth period.

### 2.3. Fungal material

Each year, we selected eight strains (**Figure 1** and **Table S1**) from a *Z. tritici* population sampled from single pycnidium on cv. Cellule and Apache in pure-stand field trials in Grignon (France) in July 2017, July 2018, and June 2019 (Orellana-Torrejon *et al*., 2022a). The virulence status of these strains with respect to *Stb16q* was determined by phenotyping on wheat cv. Cellule seedlings. The sexual compatibility of the different pairs of strains was determined by PCR amplification of the two mating-type idiomorphs (Waalwijk *et al*., 2002). Subcultures were grown for five days in Petri dishes containing PDA (potato dextrose agar, 39 g·L^−1^) at 18 °C in the dark. Blastospore suspensions of each strain were prepared as described by Suffert *et al*. (2013). The inoculum concentration was adjusted to 5 × 10^5^ conidia·mL^-1^ with a Malassez counting chamber, and two drops of surfactant (Tween 20; Sigma, France) were added. Biparental suspensions containing a Mat1.1 strain and a Mat1.2 strain were prepared by mixing 15 mL of each of the monoparental suspensions for the equiproportional combination (0.5-0.5). For the non-equiproportional combinations (unbalanced proportions: 0.1-0.9, 0.25-0.75, 0.75-0.25, and 0.9-0.1), the appropriate volumes of each suspension were combined to achieve a total volume of 30 mL.

### 2.4. Inoculation procedure

An atomiser (Ecospray, VWR, France) was used to apply each blastospore suspension onto three adult plants (nine stems) for biparental suspensions and two adult plants (six stems) for monoparental suspensions, after the wheat heads had fully emerged (on 27^th^ April 2018, 30^th^ April 2019, and 19^th^ April 2020), as described by Suffert *et al*. (2013). Plants were turned during the 10-second spraying event, to ensure even coverage with inoculum. Infection was promoted by enclosing each duo or trio of plants in a transparent polyethylene bag containing a small amount of distilled water for 72 h. The wheat plants were kept in a greenhouse for about 11-12 weeks until they had completely dried out (from 27^th^ April to 12^th^ July 2018, 30^th^ April to 9^th^ July 2019, and 19^th^ April to 8^th^ July 2020). Air temperature was kept above 24 °C during this period. In 2020, three adult plants each of Apache and Cellule were used as non-inoculated control plants.

### 2.5. Assessment of the intensity of asexual multiplication

We estimated the intensity of asexual multiplication, by determining the percentage of the leaf area covered by pycnidia (1, 2, 3 and 5%, and increments of 5% thereafter up to 100%; Suffert *et al.,* 2013) for the two uppermost leaves of each stem of the inoculated plants, between five and six weeks after inoculation (on 15^th^ June 2018, 5^th^ June 2019, and 29^th^ May 2020). Mean disease severity was, thus, calculated with 18 replicates for co-inoculations and 12 replicates for single inoculations. In the third year, an additional assessment was performed eight weeks after inoculation (on 8^th^ June 2020) on some of the leaves of cv. Cellule (co-)inoculated with avirulent strains, on which some sporulating lesions appeared very late (a few days before leaf senescence). The 17 strains collected from pycnidia in these lesions were genotyped, for comparison with the parental strains used in the experiment.

### 2.6. Promotion of sexual reproduction

After the stems and leaves had completely dried out, the wheat plants were placed outdoors during the summer and autumn, to induce the formation of pseudothecia and promote ascosporogenesis. In 2018, each duo or trio of plants was tied up with stiff wire, with the stems and leaves clamped, and placed about 10 cm away from the nearest neighbouring duo or trio, to reduce contact with these other groups of plants (from 12^th^ July 2018 to 16^th^ January 2019; ‘arrangement A’ in **Figure 2A**). In 2019 and 2020, the plants of each duo or trio were cut and grouped into small bundles, hung on a fence and spaced about 30 cm apart to ensure that contact between bundles was totally prevented (from 9^th^ July 2019 to 21^st^ January 2020 and from 22^nd^ July 2020 to 21^st^ December 2021; ‘arrangement B’ in **Figure 2B**). Each duo or trio of plants was considered to constitute a ‘set of residues’. After several weeks of exposure to outdoor weather conditions, the senescent leaves and stems from each set of residues were cut into 2 cm segments and left to dry in a laboratory atmosphere at 18 °C for one week.

**Figure 2.**
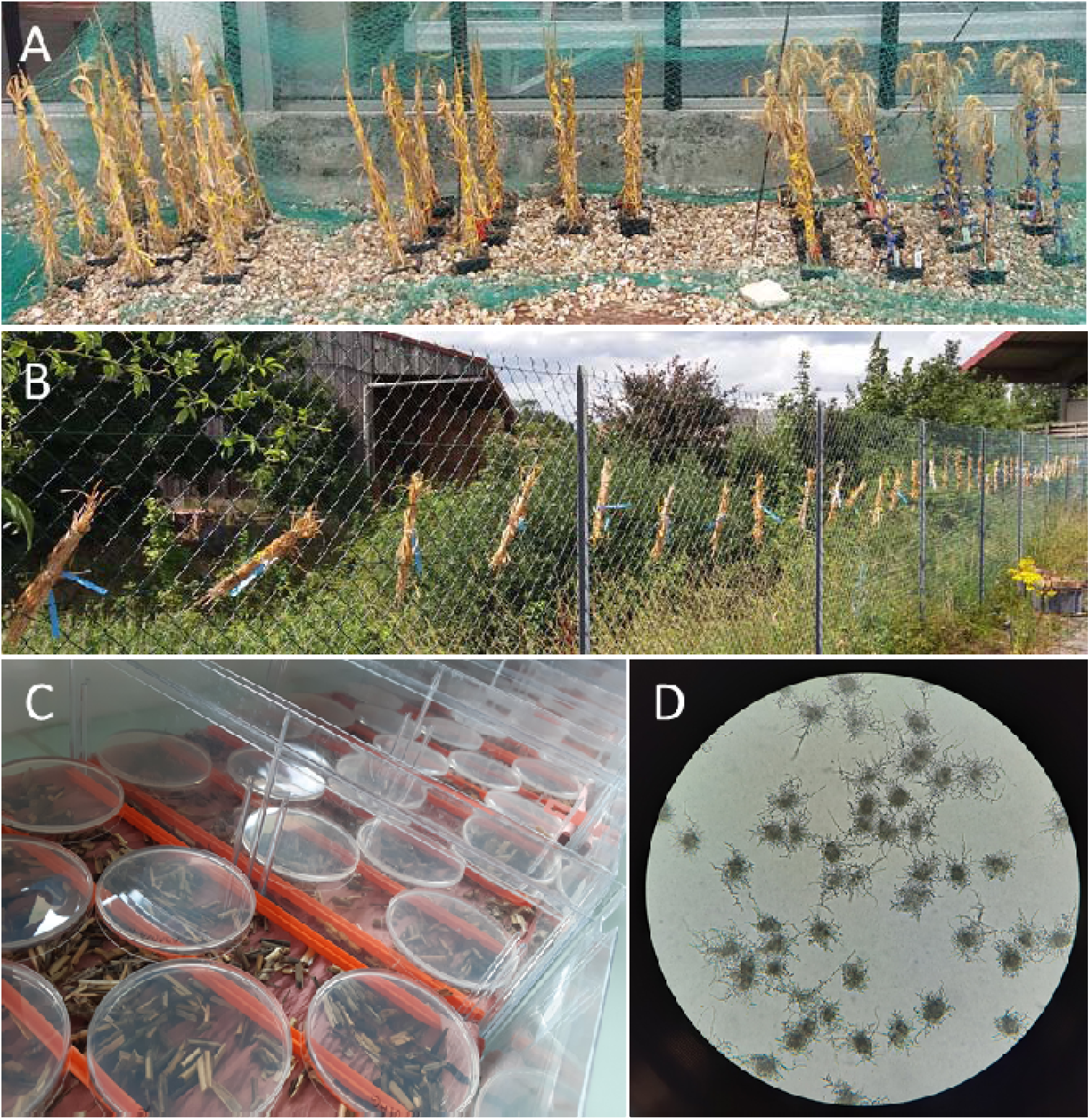
Management of sets of wheat residues, from the induction of *Zymoseptoria tritici* pseudothecia to the assessment of sexual reproduction intensity. **A**. Dry wheat plants placed outdoors (2018; ‘arrangement A’), which led to the residues partially falling on each other. **B**. Bundles of dry wheat plants hung on a fence (2019, 2020; ‘arrangement B’), with no possibility of direct contact. **C**. Petri dishes containing PDA medium placed upside down above wheat residues to collect *Z. tritici* ascospores. **D**. Ascospore-derived colonies on PDA at the time of counting, i.e., four days after discharge.

### 2.7. Assessment of the intensity of sexual reproduction

We estimated the intensity of sexual reproduction by evaluating ascospore discharge events in residue samples split in two series (16^th^ and 17^th^ January 2019, 22^nd^ and 23^rd^ January 2020, and 8^th^ and 14^th^ January 2021). Each set of residues was weighed, soaked in water for 20 min and spread on dry filter paper in a box (24 × 36 cm), the lid of which was left half-open to decrease the relative humidity of the air around the residues. Eight Petri dishes (90 mm in diameter) containing PDA medium were placed upside down 1 cm above the residues and left in the dark at 18 °C for 18 h (**Figure 2C**). The dishes were then closed and incubated in the same conditions for five days. The number of ascospore-derived colonies (**Figure 2D** **and S1**) was estimated by eye to calculate a cumulative ascospore discharge index (ADI; ascospores·g^-1^), defined as the total number of ascospores discharged per gramme of wheat residues, which we used as a proxy for sexual reproduction intensity (Suffert *et al*., 2018a, 2016, 2011).

### 2.8. Genotyping

We sampled 278 offspring strains from different ascospore-derived colonies obtained from nine single inoculations (±10 strains sampled for each) and 11 co-inoculations (±20 strains sampled for each) in 2018, 2019 and 2020. The parenthood of the offspring strains was established by genotyping the 278 offspring strains and the 24 strains used for (co-)inoculation and assumed to be their parents (**Table S1**). We used 12 SSR markers (AC0002-ST4, AG0003-ST3A, CT0004-ST9, TCC0009-ST6, AC0001-ST7, GCA0003-ST3C, chr_02-140-ST2, GAC0002-ST1, GGC0001-ST5, CAA0003-ST10, TCC0002-ST12, CAA0005-ST3B) from the panel developed by Gautier *et al*., (2014). The genotyping profiles of the offspring and parental strains were analysed with Peak Scanner software v2.0 (Applied Biosystems). We then compared these profiles, using a decision tree designed specifically for this purpose (**Figure S2**). The actual parental strains could be the strains used to inoculate the corresponding plants, other strains used to inoculate other plants during the experiment (evidence of a cross with exogenous strains from neighbouring sets of residues; ‘exo-IN’ in **Table 2**), or other strains not used in the experiment (evidence of a cross with exogenous strains from outside the experiment; ‘exo-OUT’ in **Table 2**). If all 12 SSR markers for a strain matched those of the strains used for inoculation, we considered those strains to be the true parents. However, as several of the 12 alleles are very common in natural *Z. tritici* populations, the probability of one of the parents actually being exogenous was extremely low but not zero. By contrast, if at least one of the markers in an offspring strain did not match the strains used for inoculation, we could be sure that one of the actual parents was exogenous. The ‘degree of exogeneity’ of the actual parental strains was estimated by considering the status of all genotyped offspring sampled for a given (co-)inoculation. This indicator is expressed as the percentage of offspring strains with at least one parent different from the strains used for inoculation.

We also genotyped 11 strains sampled from late asexual lesions observed on Cellule (pycnidiospores) after ‘avr × avr’ co-inoculations and six observed after single inoculations with different avirulent strains, to identify the strains responsible for these lesions.

### 2.9. Statistical analysis

All data analyses were performed with R software (v4.0.2 R Core Team 2012). The variables used to assess the intensity of asexual and sexual reproduction were non-normally distributed, as revealed by Shapiro-Wilk tests (‘shapiro.test’ function). We analysed the results of co-inoculations separately from those of single inoculations, due to a difference in the number of replicates. We, therefore, performed Kruskal-Wallis tests (*agricolae* R package, ‘kruskal’ function) for all statistical analyses concerning co-inoculations, and Wilcoxon tests (*rstatix* R package, ‘wilcox_test’ function) for all statistical analyses concerning single inoculations. Both tests were followed, when necessary, by Bonferroni corrections for pairwise comparisons, due to the high stringency of this method (*P* value threshold at 0.05). We estimated the impact on disease severity and ADI of the type of co-inoculation (‘vir × vir’, ‘avr × avr’ and ‘avr × vir’) and virulence status, for single inoculations. To this end, we first separately assessed the effects of year (2018, 2019, 2020) and cultivar (Apache, Cellule) on the two aforementioned variables, and checked for an absence of potential two-way interactions between factors. We used Benjamini & Hochberg correction for pairwise comparisons to prevent the overestimation of mean differences. For ADI, we split the data into two groups (‘arrangement A’ in 2018; ‘arrangement B’ in 2019 and 2020). We also compared the effects of single and co-inoculations on ADI and assessed the effect of the proportion of the virulent strain in the biparental ‘avr × vir’ suspensions (0.1, 0.25, 0.5, 0.75 and 0.9) on ADI and disease severity.

We used a linear model to analyse the potential correlation between the intensity of asexual multiplication and the intensity of sexual reproduction. The variance of disease severity was partitioned into sources attributable to the following factors and their interactions: ADI, the cultivar inoculated (Apache, Cellule), the type of co-inoculation and year. The model was fitted by removing non-significant interactions and we used the adjusted R² to evaluate the state of the correlation.

## 3. Results

### 3.1. Intensity of asexual multiplication resulting from single inoculations and co-inoculations

On Apache, all single inoculations caused symptoms (**Figure S3**), with significant differences in mean disease severity between strains (*p* < 0.001, Kruskall-Wallis test, data not shown). Disease severity per type of co-inoculation was scored 35 to 80% in 2018 and 2019 and 30 to 50% in 2020 (**Figure 3** and **Table S2**).

**Figure 3.**
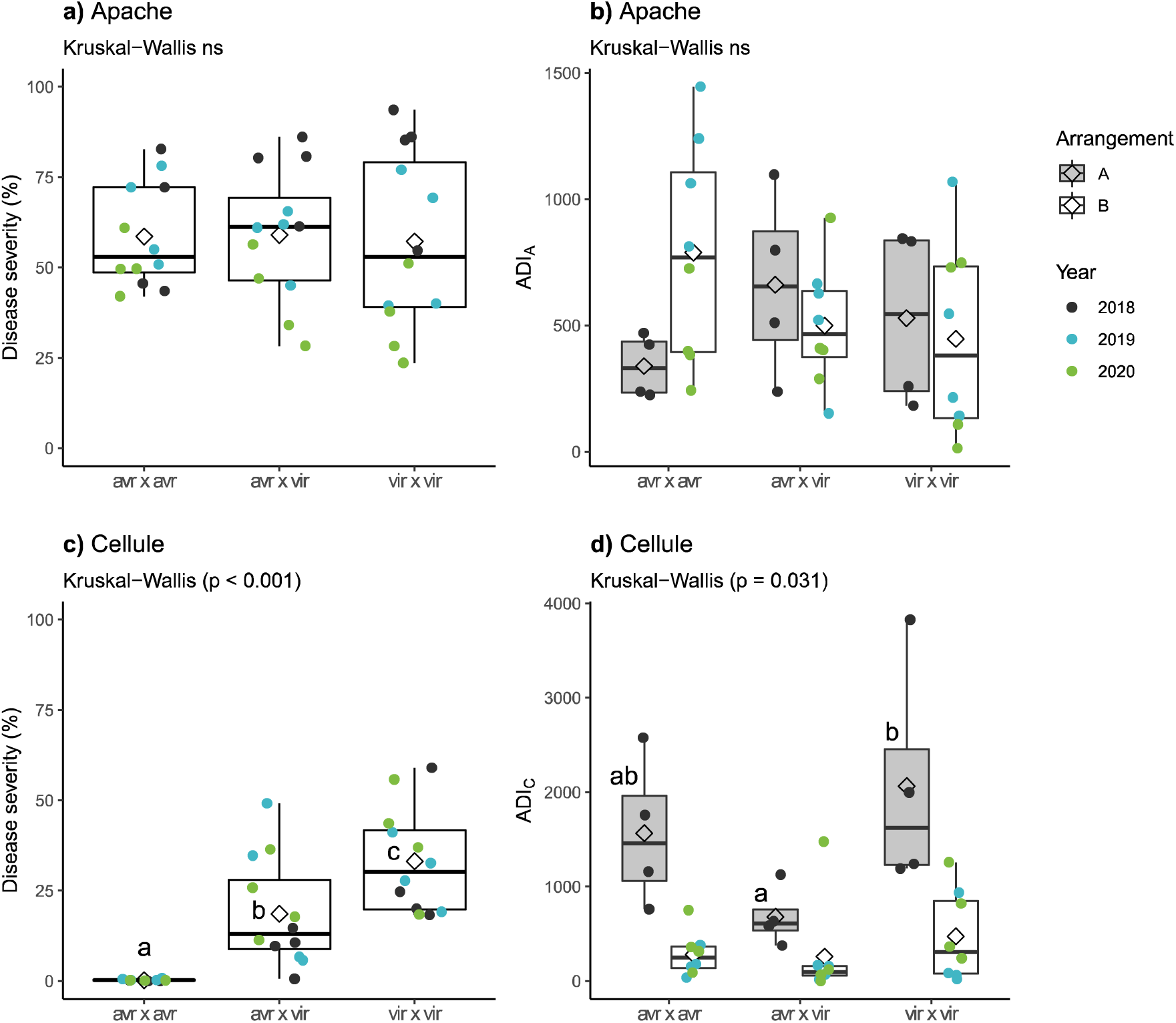
Impact of virulence status with respect to *Stb16q* of the *Zymoseptoria tritici* strains used for co-inoculation (‘avr × avr’, ‘avr × vir’, ‘vir × vir’) on the intensity of asexual multiplication (expressed as disease severity) and the intensity of sexual reproduction (expressed as the number of ascospores discharged per gramme of residues; ADI) by cultivar (Apache, Cellule) and the arrangement of the different sets of residues (‘arrangement A’ in Figure 2A; ‘arrangement B’ in Figure 2B) during the three-year experiment. Kruskal-Wallis tests, with Bonferroni correction for pairwise comparisons, were performed to compare the types of co-inoculation by cultivar and arrangement.

On Cellule, only single inoculations of virulent strains caused symptoms (**Figure S3**). We observed only rare lesions caused by single inoculations of avirulent strains (**Table S2**). Disease severity measured after ‘vir × vir’ and ‘vir × avr’ co-inoculations on Cellule was rated from 9% to 64% (**Table S2**) and was, on average, lower than that on Apache (*p* < 0.001, Wilcoxon test). As expected, no disease was observed on Cellule after ‘avr × avr’ co-inoculations (**Figure 3** and **Figure S3**).

Mean disease severity on Apache and Cellule differed significantly between years (*p* < 0.01 for both single inoculations and co-inoculations, Kruskal-Wallis test), which can be explained by the difference in disease-scoring dates between years.

Unexpectedly, some rare lesions (isolated sporulating lesions) were observed late on Cellule in 2020, eight weeks after inoculation, i.e., long after the main disease assessment (**Figure 4**). These lesions appeared in single inoculations with three different avirulent strains (FS6196, FS6503 and FS6191) and after ‘avr × avr’ co-inoculation. The genotyping of strains sampled from these asexual lesions (pycnidiospores) showed that almost all (16/17) had the same genotyping profile as one of the strains used to inoculate the plant (**Table 1 and S3**). The remaining strain had the same genotyping profile as another strain used in the experiment (FS6181) and might have resulted from of a unique contamination event in the greenhouse.

**Figure 4.**
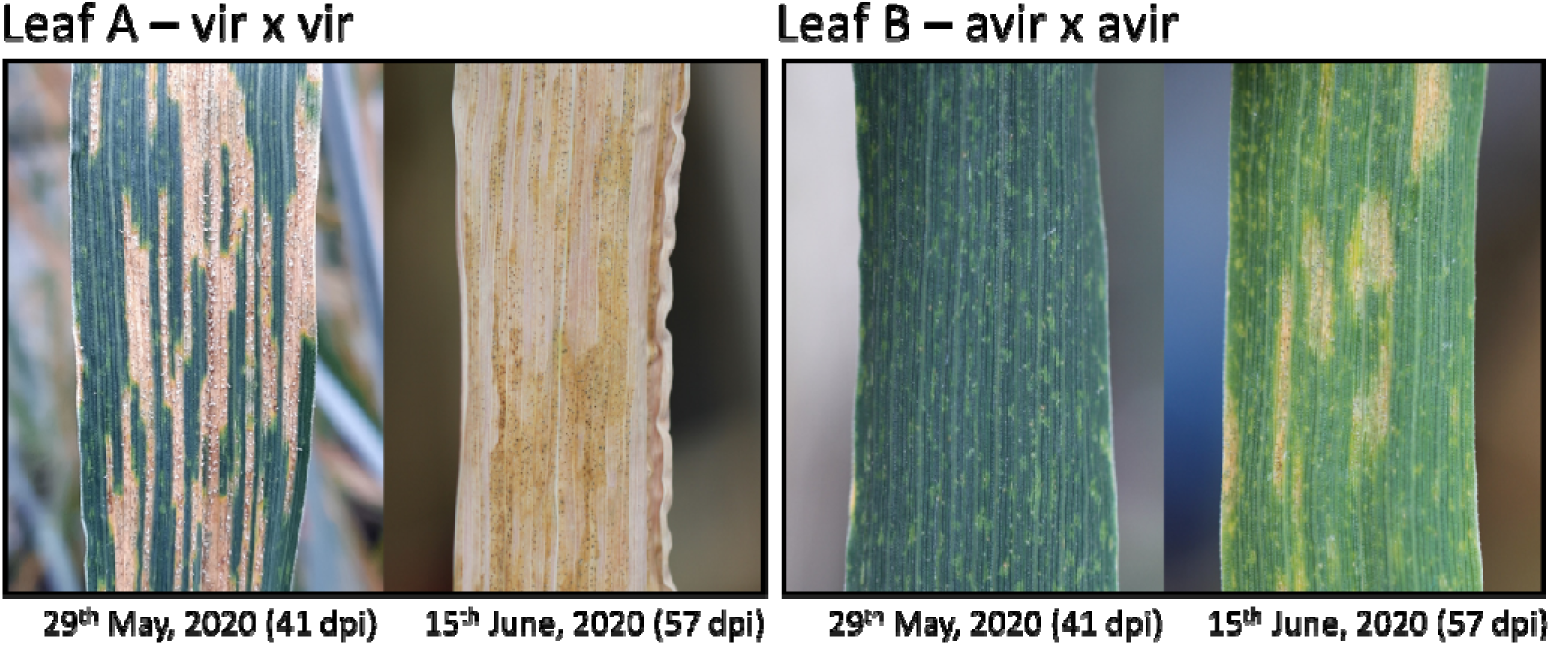
Contrasted timing of lesion appearance on adult wheat plants of cv. Cellule in 2020: five vs. eight weeks after co-inoculation. Flag leaves of Cellule were co-inoculated with either two virulent strains (Leaf A, FS6503 × FS619) or with two avirulent strains (Leaf B, FS6176 × FS6191).

**Table 1.** Identification of 17 *Zymoseptoria tritici* strains causing late lesions on wheat plants of cv. Cellule, 8 weeks after (co-)inoculation (June 15^th^, 2020), based on the comparison of their genotyping profiles with those of the 8 strains used for co-inoculation. The genotyping profiles, established with 12 SSR markers from the panel developed by Gautier *et al*. (2014), are presented in Table S3. Sampled strains that did not match the strains used for (co-)inoculation are indicated by *.

| Cv. | Strains used for<br>(co-)inoculation |  | Number of<br>strains sampled | Identity of strains<br>sampled |
| --- | --- | --- | --- | --- |
| Cellule | FS6503<br>(avr<sup>1</sup>) | - | 1 | FS6181 (× 1) * |
| Cellule | FS6191 (avr) | - | 4 | FS6191 (× 4) |
| Cellule | FS6196 (avr) | - | 1 | FS6196 (× 1) |
| Cellule | FS6503 (avr) | FS6191 (avr) | 3 | FS6191 (× 3) |
| Cellule | FS6503 (avr) | FS6196 (avr) | 3 | FS6196 (× 3) |
| Cellule | FS6522 (avr) | FS6191 (avr) | 2 | FS6191 (× 2) |
| Cellule | FS6522 (avr) | FS6196 (avr) | 3 | FS6196 (× 3) |

<sup>1</sup> 'avr' for avirulent against *Stb16q*

### 3.2. Intensity of sexual reproduction and genotyping of offspring resulting from co-inoculations

Overall, the intensity of sexual reproduction depended on cultivar in 2018 and 2019 (*p* = 0.002, Wilcoxon), on year for Cellule (*p* < 0.001, Kruskal-Wallis test; **Figure 5**), and on the type of co-inoculation, as described below for each cultivar.

**Figure 5.**
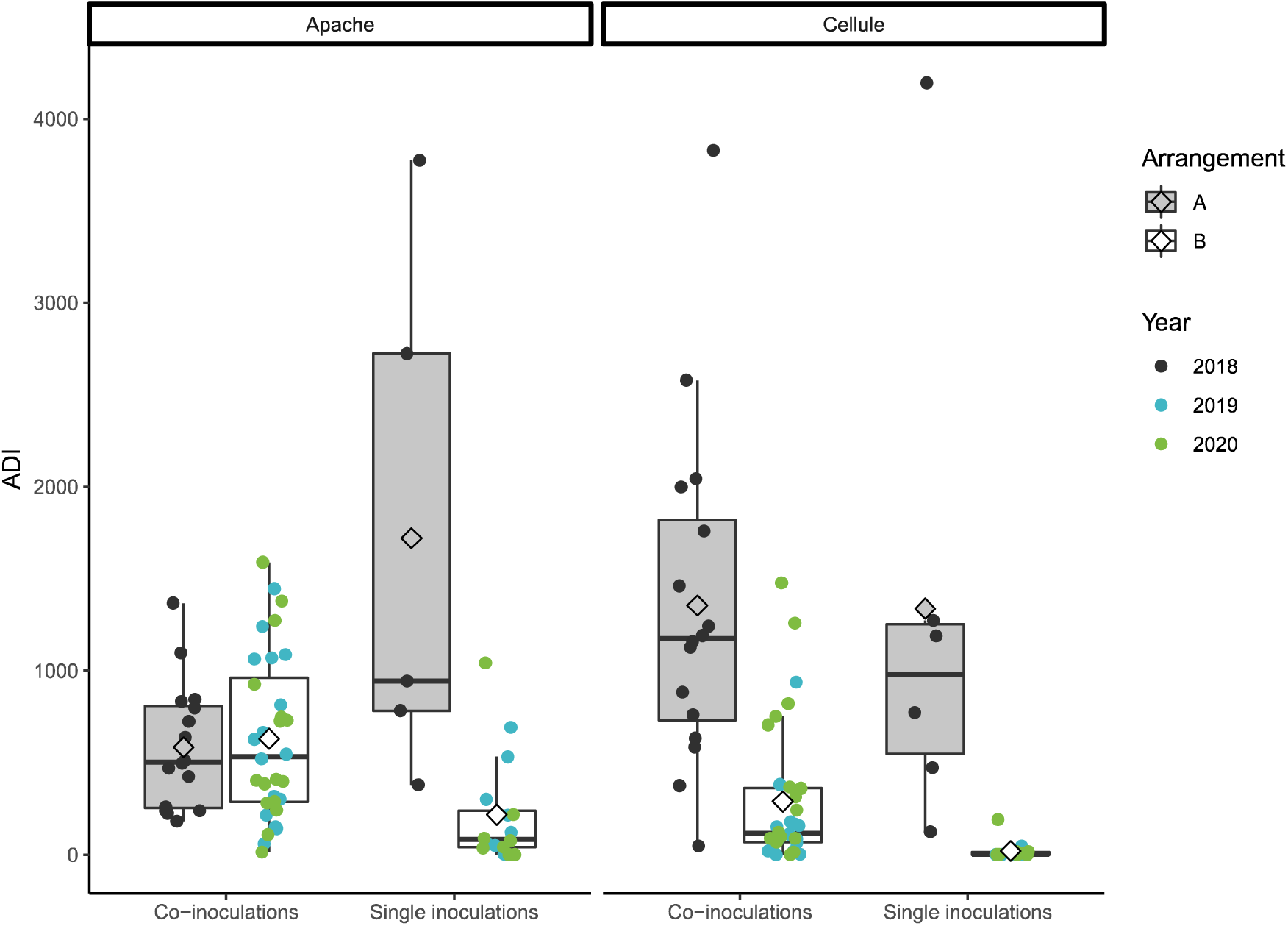
Impact of the type of inoculation (single inoculations, co-inoculations) on the intensity of *Zymoseptoria tritici* sexual reproduction (expressed as the number of ascospores discharged per gramme of residues; ADI), by cultivar (Apache, Cellule) and the arrangement of the different sets of residues (‘arrangement A’ in Figure 2A; ‘arrangement B’ in Figure 2B) during the three-year experiment. Kruskal-Wallis tests, with Bonferroni correction for pairwise comparisons, were performed to compare the types of co-inoculation by cultivar and arrangement.

On Apache, we obtained offspring strains for all type of co-inoculations (‘vir × vir’, ‘avr × avr’ and ‘avr × vir’) as expected, regardless of the virulence or avirulence status of parental strains (**Figure 5** and **Table S4**). The intensity of sexual reproduction on this cultivar was not dependent on the virulence status of the parental strains: there were no significant differences in mean ascospore discharge index (ADI) between the three types of co-inoculation (**Figure 5** and **Table S4**). This is not surprising, as the virulence status of strains is relevant only for Cellule, the cultivar carrying the corresponding resistance gene.

On Cellule, we obtained offspring strains for ‘vir × vir’ co-inoculations, as expected, but also for ‘vir × avr’ and even for ‘avr × avr’ co-inoculations (**Figure 5**). The intensity of sexual reproduction depended on the virulence status of the parental strains: ADI was higher for ‘vir × vir’ than for ‘vir ×x avr’ and ‘avr ×x avr’ inoculations, with significant differences only in 2018 (*p* = 0.031, Kruskal-Wallis; **Figure 5** and **Table S4**).

The analysis of genotyping data for offspring strains sampled from Cellule residues confirmed that the parental strains actually involved in sexual reproduction were very probably the strains used for inoculation (**Table 2, Table S3** and **Figure S2A**). Few cases were identified in which at least one of the actual parental strains was not among those used for inoculation (mean degree of exogeneity of the parental strains: 5% for all co-inoculations). The genotyping results demonstrate that an avirulent strain present asymptomatically on a plant (without causing sporulating lesions five weeks after inoculation) could cross with a virulent strain (infecting). For instance, two ‘vir × avr’ co-inoculations, FS2883 × FS2847 in 2018 and FS6181 × FS6196 in 2020, produced offspring for which we genotyped a sample (20 and 25 strains, respectively): all alleles of the 12 SSR markers for all offspring strains matched those of the parental strains used for co-inoculation (no exogeneity of the parental strains; **Table 2**). Moreover, genotyping demonstrated that two avirulent strains present asymptomatically on a plant — at least five to six weeks after inoculation — were also able to cross. This is the most striking finding of this study. The ‘avr × avr’ co-inoculations, FS2851 × FS2855 and FS2849 × FS2847 in 2019 and FS6503 × FS6196 and FS6522 × FS6196 in 2020 all produced offspring strains in which all alleles of the 12 SSR markers matched those of the parental strains used for co-inoculation (no exogeneity of the parental strains; **Table 2**). The genotyping results for the offspring strains from the ‘vir × avr’ co-inoculations FS1492 × FS1721 and FS1425 × FS1754 in 2018 and the ‘avr × avr’ co-inoculation FS1721 × FS1754 are consistent with the results reported above, even though the degree of exogeneity of the parental strains was non-null (mean value of 16%; **Table 2**). Four of the 51 offspring strains genotyped resulted from a cross between one of the parents used for co-inoculation and another parental strain used in the experiment (‘exo-IN’), and three resulted from a cross between one of the parental strains used for co-inoculation and a strain from outside the experiment (‘exo-OUT’; **Table 2**). The parental strains displayed a higher degree of exogeneity in 2018 (**Table 2**), probably due to the difference in the arrangement of the residues between this and the other two years (**Figure 2A** **and 2B**).

**Table 2.**
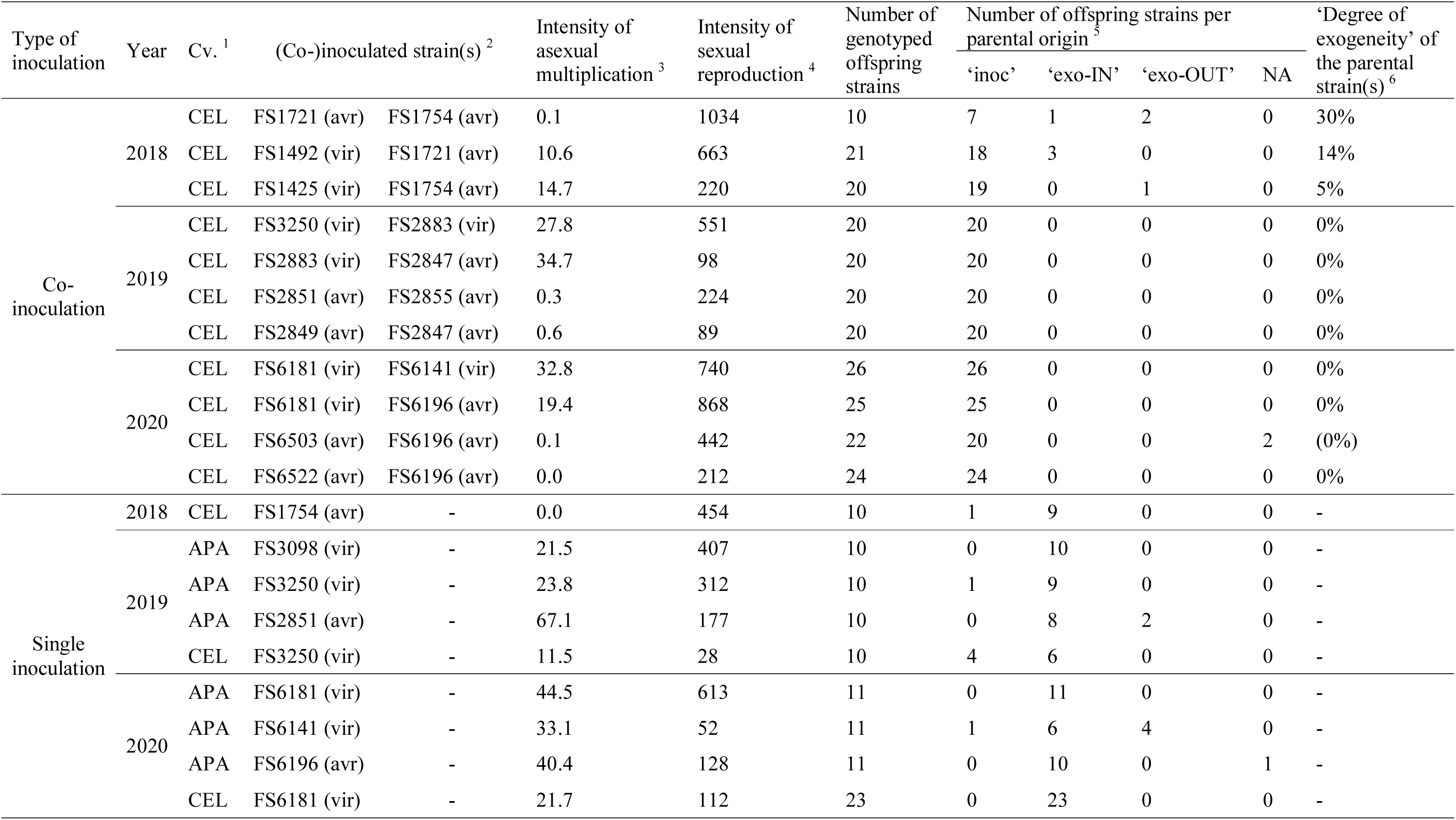
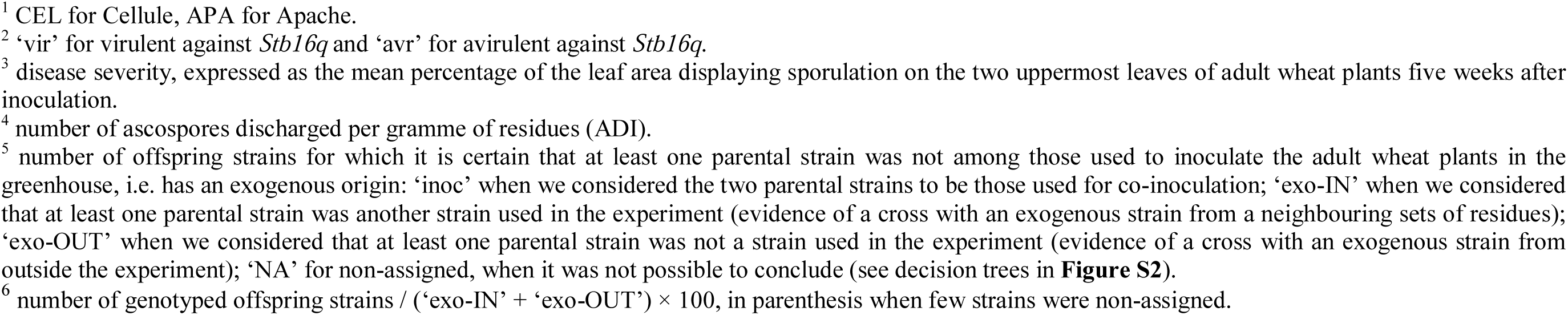
Parenthood of 278 *Zymoseptoria tritici* offspring strains sampled from ascospore-derived colonies obtained after nine single inoculations and 11 co-inoculations (a subset of all the inoculations performed in the three-year experiment). Parenthood was established by comparing the genotypic profiles of the 278 strains with those of the three sets of eight strains used for (co-)inoculation, based on the decision tree presented in **Figure S2**. The genotypic profiles, established with 12 SSR markers from the panel developed by Gautier *et al*. (2014), are presented in Table S3.

| Type of inoculation | Year | Cv. <sup>1</sup> | (Co-)inoculated strain(s) <sup>2</sup> |  | Intensity of asexual multiplication <sup>3</sup> | Intensity of sexual reproduction <sup>4</sup> | Number of genotyped offspring strains | Number of offspring strains per parental origin <sup>5</sup> |  |  |  | 'Degree of exogeneity' of the parental strain(s) <sup>6</sup> |
| --- | --- | --- | --- | --- | --- | --- | --- | --- | --- | --- | --- | --- |
|  |  |  |  |  |  |  |  | 'inoc' | 'exo-IN' | 'exo-OUT' | NA |  |
| Co-inoculation | 2018 | CEL | FS1721 (avr) | FS1754 (avr) | 0.1 | 1034 | 10 | 7 | 1 | 2 | 0 | 30% |
|  |  | CEL | FS1492 (vir) | FS1721 (avr) | 10.6 | 663 | 21 | 18 | 3 | 0 | 0 | 14% |
|  |  | CEL | FS1425 (vir) | FS1754 (avr) | 14.7 | 220 | 20 | 19 | 0 | 1 | 0 | 5% |
|  | 2019 | CEL | FS3250 (vir) | FS2883 (vir) | 27.8 | 551 | 20 | 20 | 0 | 0 | 0 | 0% |
|  |  | CEL | FS2883 (vir) | FS2847 (avr) | 34.7 | 98 | 20 | 20 | 0 | 0 | 0 | 0% |
|  |  | CEL | FS2851 (avr) | FS2855 (avr) | 0.3 | 224 | 20 | 20 | 0 | 0 | 0 | 0% |
|  |  | CEL | FS2849 (avr) | FS2847 (avr) | 0.6 | 89 | 20 | 20 | 0 | 0 | 0 | 0% |
|  | 2020 | CEL | FS6181 (vir) | FS6141 (vir) | 32.8 | 740 | 26 | 26 | 0 | 0 | 0 | 0% |
|  |  | CEL | FS6181 (vir) | FS6196 (avr) | 19.4 | 868 | 25 | 25 | 0 | 0 | 0 | 0% |
|  |  | CEL | FS6503 (avr) | FS6196 (avr) | 0.1 | 442 | 22 | 20 | 0 | 0 | 2 | (0%) |
|  |  | CEL | FS6522 (avr) | FS6196 (avr) | 0.0 | 212 | 24 | 24 | 0 | 0 | 0 | 0% |
| Single inoculation | 2018 | CEL | FS1754 (avr) | - | 0.0 | 454 | 10 | 1 | 9 | 0 | 0 | - |
|  | 2019 | APA | FS3098 (vir) | - | 21.5 | 407 | 10 | 0 | 10 | 0 | 0 | - |
|  |  | APA | FS3250 (vir) | - | 23.8 | 312 | 10 | 1 | 9 | 0 | 0 | - |
|  |  | APA | FS2851 (avr) | - | 67.1 | 177 | 10 | 0 | 8 | 2 | 0 | - |
|  |  | CEL | FS3250 (vir) | - | 11.5 | 28 | 10 | 4 | 6 | 0 | 0 | - |
|  | 2020 | APA | FS6181 (vir) | - | 44.5 | 613 | 11 | 0 | 11 | 0 | 0 | - |
|  |  | APA | FS6141 (vir) | - | 33.1 | 52 | 11 | 1 | 6 | 4 | 0 | - |
|  |  | APA | FS6196 (avr) | - | 40.4 | 128 | 11 | 0 | 10 | 0 | 1 | - |
|  |  | CEL | FS6181 (vir) | - | 21.7 | 112 | 23 | 0 | 23 | 0 | 0 | - |

<sup>1</sup> CEL for Cellule, APA for Apache.
<sup>2</sup> ‘vir’ for virulent against *Stb16q* and ‘avr’ for avirulent against *Stb16q*.
<sup>3</sup> disease severity, expressed as the mean percentage of the leaf area displaying sporulation on the two uppermost leaves of adult wheat plants five weeks after inoculation.
<sup>4</sup> number of ascospores discharged per gramme of residues (ADI).
<sup>5</sup> number of offspring strains for which it is certain that at least one parental strain was not among those used to inoculate the adult wheat plants in the greenhouse, i.e. has an exogenous origin: ‘inoc’ when we considered the two parental strains to be those used for co-inoculation; ‘exo-IN’ when we considered that at least one parental strain was another strain used in the experiment (evidence of a cross with an exogenous strain from a neighbouring sets of residues); ‘exo-OUT’ when we considered that at least one parental strain was not a strain used in the experiment (evidence of a cross with an exogenous strain from outside the experiment); ‘NA’ for non-assigned, when it was not possible to conclude (see decision trees in **Figure S2**).
<sup>6</sup> number of genotyped offspring strains / (‘exo-IN’ + ‘exo-OUT’) × 100, in parenthesis when few strains were non-assigned.

### 3.3. Intensity of sexual reproduction and genotyping of offspring resulting from single inoculations

Another key finding of this study was that offspring strains could be obtained following single inoculations, whereas plants not inoculated in 2020 (controls) produced no offspring. At first sight, this result is surprising, as it shows that sexual reproduction does not necessarily require inoculation with the two parental strains such that they are present in or on (growing epiphytically) the host tissues whilst the plant is growing. Surprising as it may seem, this result was obtained in each of the three years (**Table 2,** **Figure 5**). The “unexpected” crosses can be explained by the arrival of “exogenous” strains on residues (after the senescent plants were placed outside) from neighbouring sets of residues for the strains used in the experiment (‘exo-IN’) and from residues present in fields away from the experimental site (‘exo-OUT’). In 2018, most of the offspring strains obtained after the inoculation of Cellule with a single avirulent strain (FS1754) were considered to result from a cross between the strain used for inoculation and another parental strain used in the experiment (**Table S3** and **Figure S2A**). This can be explained by the difference in the arrangement of the residues (A vs. B; **Figure 2**). This result is particularly surprising given that the avirulent parental strain used for inoculation produced no symptoms on the plant during the asexual stage. In 2019 and 2020, despite the distancing of residues, offspring strains were obtained after single inoculations, whatever the cultivar or the virulence status of the strain used for inoculation. We genotyped 42 and 21 offspring strains sampled from residues of Apache after single inoculation with a virulent or avirulent strain, respectively. We found that 36 (86%) and 18 (86%) offspring strains, respectively, could be considered to result from a cross between the strain used for inoculation and another parental strain used in the experiment (‘exo-IN’), and that 4 (10%) and 2 (10%), respectively, resulted from a cross between the inoculated strain and strain from outside the experiment (‘exo-OUT’; **Table 2**, **Figure S2B**). For the remaining cases (two and one, respectively), the genotyping profiles of the offspring strains matched those of the strains used for inoculation, and we cannot, therefore, conclude on the origin of the exogenous parent (‘exo-IN’ or ‘exo-OUT’). On Cellule, 29 of the 33 offspring strains sampled after single inoculation with a virulent strain (88%) were considered to result from a cross between the inoculated strain and another of the parental strains used in the experiment. The remaining four strains (12%) had the same genotyping profile as the strain inoculated, and it was not, therefore, possible to draw any conclusions about the origin of the exogenous parent (**Table 2**, **Figure S2B**).

The intensity of sexual reproduction was lower after single inoculations than after co-inoculations in 2019 and 2020 (**Figure 5**). Support for the role of non-inoculated strains (arriving on host tissues after the plant was fully senescent) in successful crosses is provided by the observation that the intensity of sexual reproduction in single inoculations was dependent on year (*p* = 0.056 for avirulent strains in Apache and *p* = 0.002 in Cellule, Kruskal-Wallis test).

### 3.4. Impact of diverse factors on the intensity of sexual reproduction

#### Impact of environmental conditions

As highlighted above, the intensity of sexual reproduction was significantly dependent on year, with higher values for ‘avr × avr’ co-inoculations in Apache in 2018 (*p* = 0.023, Kruskal-Wallis) and for ‘vir × vir’ and ‘avr × avr’ co-inoculations in Cellule in 2018 than for the corresponding co-inoculations in 2019 and 2020 (*p* = 0.03, *p* = 0.02, respectively, Kruskal-Wallis; **Table S4**). This was also the case for single inoculations in Apache and Cellule in 2018 (*p* = 0.011 and *p* = 0.002, respectively, Kruskal-Wallis). The impact of year is consistent with the difference in the arrangement of the sets of residues (A vs. B): “cross-contaminations” between sets of residues were frequent in 2018, when the arrangement of residues made physical contact possible, but were more limited in 2019 and 2020, when the sets of residues were placed further apart. The intensity of sexual reproduction was higher in field conditions in 2018 than in 2019, and this may be attributed to differences in climatic conditions, as highlighted in a previous study (Orellana-Torrejon *et al*., 2022a).

#### Impact of the proportions of the parental strains

The intensity of sexual reproduction (ADI) was not significantly affected by the proportion (0.1, 0.25, 0.5, 0.75, and 0.9) of the virulent parental strain in the three pairs of ‘avr × vir’ strains tested, whatever the wheat cultivar, probably because of the small number of replicates and the high variability of ADI (**Table S5**). ADI was higher on Apache than on Cellule (*p* = 0.025 Wilcoxon test), with no significant difference between years in any cultivar, and it tended to be higher for Cellule in 2018 (*p* = 0.086, Kruskal-Wallis test), when ‘arrangement A’ was applied. By grouping similar unbalanced proportions (group 1/10, combining the 0.1-0.9 and 0.9-0.1; group 1/4 combining 0.25-0.75 and 0.75-0.25), we found that the intensity of sexual reproduction after co-inoculation in unbalanced proportions was higher than that when parental strains were used in equal proportions for co-inoculation (0.5-0.5), whatever the cultivar. As expected, disease severity on Apache did not differ with the proportion of the virulent parental strain (**Table S6**), whereas disease severity increased significantly with the proportion of the virulent strain on Cellule.

## 4. Discussion

### 4.1. “Unexpected” crosses of avirulent strains in plants carrying S*tb16q*: from experimental findings to explanatory hypotheses

We showed that sexual reproduction between virulent (‘vir’) and avirulent (‘avr’) strains is possible in a wheat cultivar carrying the corresponding resistance gene, here *Stb16q* (process ② in **Figure 6**; see also **Figure 5** **and S3**, **Table 2**). This finding is consistent with the results obtained by Kema *et al*. (2018) in a similar experiment in which all *Z. tritici* crosses on seedlings of a cultivar carrying *Stb6* were successful even if one of the parental strains was avirulent, with no pathogenic relationship to the wheat cultivar used. More unexpectedly, we found that two avirulent strains could cross without causing symptoms in plants, five weeks after their use for co-inoculation (see below). For several crossing situations (**Table 2** and **Figure S2**), we established by genotyping that the strains used for co-inoculation could be considered to be the actual parents, even if the host-pathogen interaction during the asexual stage was “incompatible”. As a means of understanding the mechanisms underlying the ‘vir’× ‘avr’ and ‘avr × avr’ crosses, we focused on the rare symptoms (**Figure 4B**) observed on Cellule late after inoculation in 2020, with an origin — the strains used for inoculation — confirmed by genotyping (**Table 1**). The immunity conferred by *Stb16q* throughout growth of the wheat plants prevented complete infection until the appearance of lesions by limiting pathogen progression in the substomatal cavities or the apoplast (Saintenac *et al*., 2021). We hypothesize that this type of immunity ceased to operate just before plant senescence, allowing the pathogen to colonise the host tissues more deeply in some areas of the leaf and leading to the appearance of a few isolated lesions (**Figure 4D**). This observation of late symptoms is also consistent with the epiphytic development and/or survival of *Z. tritici* on the surface of leaves, as reported by Fones *et al*. (2017). This epiphytic development may be explained by specific experimental conditions in the greenhouse: application of a high concentration of inoculum on the leaf surface and favourable conditions for fungal growth over an extended period of time. This phenomenon could however happen in the field: 3.3% of the strains sampled on Cellule grown in mixture with Apache, from STB lesions, were found to be avirulent against *Stb16q* (Orellana-Torrejon *et al*., 2022a). The success of ‘vir × avr’ crosses (process ② in **Figure 6**) can be also explained by the hypothesis that penetration of the avirulent strain was favoured by a virulent strain that had previously infected the host tissues (Tollenaere *et al*., 2016). Facilitation mechanisms of this type have been reported in multiple infections in some pathosystems. For example, the initial arrival of a virulent strain of *Blumeria graminis* on a host plant can suppress resistance, enabling subsequently arriving avirulent strains to penetrate host tissues (Lyngkjær *et al*., 2001). The possibility of infection by “stowaway” strains through a ‘systemic induced susceptibility’ reaction was experimentally proposed for *Z. tritici* by Seybold *et al*. (2020) and recently discussed by Barrett *et al*. (2021). This hypothesis is consistent with the intensity of sexual reproduction tending to be higher for unbalanced proportions of the two parental strains, as observed in our study (see 4.4), which could promote the penetration of the avirulent strain. However, this finding requires experimental confirmation and should, therefore, be interpreted with caution.

**Figure 6.**
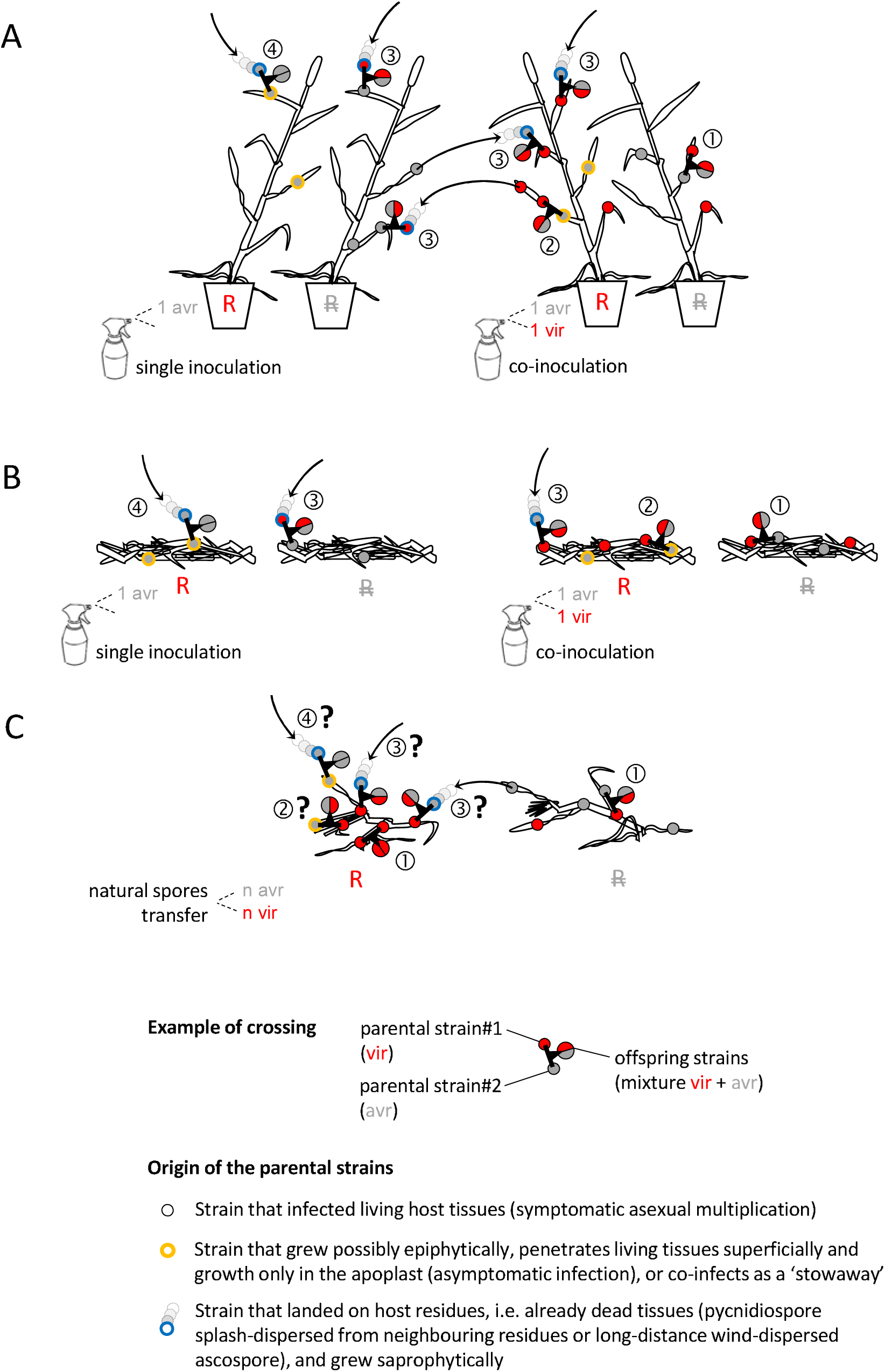
Epidemiological processes, including spore transfers, leading to sexual crosses on wheat residues according to the origin of the *Zymoseptoria tritici* parental strains and their virulence status, (**A**, **B**) in the conditions of the three-year experiment and (**C**) in field conditions. Each crossing event is characterised by cultivar — with or without a resistant gene R (Cellule R, Apache) — where the cross occurs, the virulence status (‘vir’, ‘avr’) of the parental strains, and the status of the offspring. Four crossing scenarios are proposed according to the origin and type of growth of the parental strains: ① both parents (virulent strains) infect independently living tissues; ② one parent (virulent strain, acting as the maternal parent) infects living tissues whereas the other parent (avirulent strain) grows asymptomatically, possibly epiphytically, penetrates living tissues superficially and growth only in the apoplast (asymptomatic infection), or co-infects as a ‘stowaway’ taking advantage of the infection with the other strain; ③ one parent (virulent strain) infects living tissues whereas the other parent (virulent or avirulent strain) arrives later on residues from a neighbouring set of residues (pycnidiospores; possible in **A** and **C**, but not possible in **B**) or after covering a longer distance (ascospores; possible in **A**, **B**, **C**) and then grows saprophytically; ④ one parent (avirulent strain) grows asymptomatically, possibly epiphytically, whereas the other parent (virulent or avirulent strain) arrives later on residues from a neighbouring set of residues (pycnidiospores; possible in **A** and **C**, but not possible in **B**) or after covering a longer distance (ascospores; possible in **A**, **B**, **C**) and then grows saprophytically.

### 4.2. Parental strains do not necessarily need to infect their living host to reproduce sexually: the epidemiological processes potentially involved

Debuchy *et al*. (2010) suggested that the physical proximity of parental strains might be sufficient to induce the development of reproductive organs when abiotic conditions are optimal, in a manner dependent on the pathogenic fungi concerned. The recognition of sexually compatible strains is based on the reception of pheromones for heterothallic fungi (Ni *et al*., 2011). There is, therefore, no evidence that sexual reproduction consistently requires at least one of the two parents to infect the living host tissues until the appearance of symptoms. The “unexpected” crossing situations highlighted in this study support this assumption and suggest that avirulent strains may have developed on leaves, possibly epiphytically or may have penetrated the host tissues only superficially (Fones *et al*., 2017; Fantozzi *et al*., 2021). Microscopy studies showed that avirulent strains are able to germinate and penetrate into resistant plants tissues (Cohen & Eyal, 1993; Shetty *et al*., 2003) but that the infection process is then blocked: pathogen can growth in the apoplast and plant cells are not penetrated. The resistance based on major resistance genes active against apoplastic fungal pathogens is not actually associated with fungal death (Stotz *et al*., 2014), supporting the idea that avirulent strains may survive in resistant host tissues. Saintenac *et al*. (2021) provided experimental evidence of this for *Stb16q*: they detected fungal biomass 21 days after the inoculation of a resistant cultivar with an avirulent strain of *Z. tritici*, despite the absence of symptoms due to host-pathogen incompatibility. The results obtained by Kema *et al*. (2018) for *Z. tritici,* focusing on *Stb6,* suggest that the “female” parental strain must infect the host tissues to form maternal ascogonia: the skew in the segregation of mitochondrial SSRs (and, thus, an unbalanced ratio for maternal or paternal parenthood) obtained after co-inoculations showed that the avirulent parent was almost exclusively the paternal parent. This finding is consistent with hypothetical process ③ in **Figure 6**. However, as we obtained crosses on Cellule after single inoculations with avirulent strains (not only in 2018, but also, to a lesser extent, in 2019 and 2020; **Figure 5** and **Table S4**), it remains possible that ascogonia were also formed on senescent tissue after the maternal strain had grown saprophytically (process ④ in **Figure 6**).

Our results also suggest that the second exogenous parental (probably paternal) strain arrived later, on senescent tissues, when the set of residues had been placed outdoors, and that it grew saprophytically before crossing. Sexual reproduction may, therefore, have been initiated relatively late, at a stage equivalent to the interseason period in field conditions. Furthermore, this scenario is the only one capable of explaining the crosses observed following inoculation with a single virulent or avirulent strain (processes ③ and ④, respectively, in **Figure 6**; **Figure 5** and **Table 2**). The exogenous parental strain may have come from: (i) pycnidiospores transferred from neighbouring residues by rainsplash or direct physical contact (which was possible in the 2018 arrangement), or (ii) ascospores dispersed by the wind from field residues farther away in 2019 and 2020. Further experiments are required to validate processes ② ③ ④ in addition to the classical process ① for each crossing situation. For instance, the direct inoculation of residues with the second parental strain (or both parental strains) might make it possible to demonstrate the ability of a strain to undergo sexual reproduction after growing saprophytically, as for *Phaeosphaeria nodorum* (Halama & Lacoste, 1992). Cytological analyses would be relevant, but complex to perform on senescent tissues. The current state of knowledge and hypotheses about sexual reproduction are consistent with the hemibiotrophic status of *Z. tritici* and a necrotrophic status for *P. nodorum*, although the lifestyles of these function have been defined to date on the sole basis of analyses of the asexual stage (Sanchez-Vallet *et al*., 2015).

### 4.3. Relationship between the intensity of asexual multiplication and the intensity of sexual reproduction

We found no correlation (R^2^ = 0.38) between the intensity of asexual multiplication and the intensity of sexual reproduction (**Figure S3**). Suffert *et al*. (2018a) highlighted a significant positive correlation for disease severity scores that were lower (from 10% to 40%) than those reported here (35% to 80% on Apache), despite the similar experimental conditions. In this study, disease severity levels were high enough to maximize physical contact between the two parental strains, but also to induce a decrease in the intensity of sexual reproduction (‘density-dependent’ processes), as previously shown by Suffert *et al*. (2018a).

### 4.4. Impact of the proportions of parental strains: a trend requiring confirmation

We found no significant impact of the proportions of parental strains on the intensity of sexual reproduction (**Table S4**). However, we showed, by grouping similar parental strain ratios (1/10 and 1/4), that the intensity of sexual reproduction tended to be higher for unbalanced proportions of parental strains. This trend, consistent with the increase in the intensity of sexual reproduction highlighted by Suffert *et al*. (2016) when parental strains had latent periods differing by a few days, requires confirmation in future experiments.

### 4.5. Epidemiological consequences for avirulent strains reproducing sexually on resistant cultivars: a possibility that could be exploited to limit the breakdown of a resistance gene?

By contrast to our findings, all crosses involving two avirulent parental strains failed in the study by Kema *et al*. (2018). These divergent results may be explained by differences in the plant stages considered: adult plants treated until natural senescence and with a long crossing time in our study, versus seedlings and a short crossing time in the study by Kema *et al*. (2018). Moreover, we found that the most classical scenario of ‘vir × vir’ crosses (process ① in **Figure 6**) and the “unexpected” ‘vir × avr’ and ‘avr × avr’ scenarios on plants carrying a resistance gene generated offspring populations of similar sizes. Thus, not only can avirulent strains reproduce sexually, but they can also produce substantial numbers of ascospores. This finding conflicts with the paradigm that avirulent strains tend to be eliminated from a population because they do not infect the host, and cannot therefore reproduce and contribute to subsequent epidemics, at least in agrosystems characterised by monovarietal fields. In fact, avirulence genes can be transmitted to the next generation even in a wheat canopy considered resistant. This can temporally affect the local evolution of the pathogen population and, hence, the dynamics of resistance breakdown. The extent of resistance breakdown should depend: (i) on the number of avirulent strains arriving on the cultivar carrying the resistance gene, and (ii) the available opportunities for encountering and crossing with other strains. ‘Vir × avr’ and ‘avr × avr’ crosses are probably rare in pure stands (the lower the frequency of avirulent strains in the pathogen population, the rarer), but their frequency could be enhanced by certain resistance deployment strategies. For instance, the use of cultivar mixtures would increase the exposure of plants carrying a resistance gene to the corresponding avirulent strains, due to the presence of neighbouring susceptible plants on which these strains would be then able to multiply asexually. Orellana-Torrejon *et al*. (2022a, 2022b) highlighted a consequence of this phenomenon in the field: the frequency of avirulent *Z. tritici* strains was higher after sexual reproduction on the residues of a resistant wheat cultivar grown in a mixture than after sexual reproduction on the residues of a pure stand, providing experimental evidence of an “overtransmission” of avirulence to the next epidemic season. Cultivar mixtures, therefore, appear to be a promising way to exploit the epidemiological consequences of avirulent strains crossing on resistant cultivars for limiting the breakdown of resistance genes.

## 5. Conclusion

The selection pressure exerted on pathogen populations by the deployment of a resistance gene in fields planted with a single cultivar promotes the generalisation of virulent strains (Brown and Tellier, 2011). Virulence is, theoretically, accompanied by a fitness cost (Burdon and Laine, 2019), but it was thought that this advantage for avirulent strains (rarely demonstrated; see, for instance, Orellana-Torrejon *et al*., 2022a) was compensated by their inability to persist in pathogen populations over a long period of time. The experimental findings presented here challenge partly this assumption. The hypothetical epidemiological processes we propose can explain the diversity of possible crossing scenarios (detailed in **Figure 6**) contributing to the maintenance of avirulence in *Z. tritici* populations. Epi-evolutionary models incorporating sexual reproduction should take the diversity of these scenarios into account, to improve our understanding of their impact on the transmission of avirulence in pathogen populations (Fabre *et al*., 2022; Rimbaud *et al*., 2021), particularly when the aim is to compare different deployment strategies for cultivar resistances at large spatiotemporal scales.

## Acknowledgements

We thank Marie-Pierre Guillot (AgroParisTech BIOGER, Thiverval-Grignon, FR), Nathalie Retout and Laurent Gerard (INRAE BIOGER, Thiverval-Grignon, FR) for technical assistance, Dr. Cyrille Saintenac (INRAE GDEC, Clermont-Ferrand, FR), Dr. Romain Valade (ARVALIS-Institut du Végétal, Boigneville, FR) and Dr. Marc-Henri Lebrun (INRAE BIOGER, Thiverval-Grignon, FR) for sharing information and relevant suggestions throughout this project. We also thank Dr. Julie Sappa for her help correcting our English.

## Data Availability Statement

The raw data used for the analysis of the intensity of asexual multiplication and the intensity of sexual reproduction of *Z. tritici* are available from the INRAE Dataverse online data repository (https://data.inrae.fr/) at https://doi.org/10.15454/IPMIOX.

---

**Table S1.** List of the 24 *Zymoseptoria tritici* strains used as parental strains in the three-year experiment. Strains were sampled on wheat cv. Cellule and Apache in pure-stand field trials located in Grignon (France) in July 2017, July 2018, and June 2019 (Orellana-Torrejon et al., 2022a).

| Year | Code | Mating type <sup>1</sup> | Wheat cv.<br>of origin | Virulence status <sup>2</sup> |
| --- | --- | --- | --- | --- |
| 2018 | (INRA17-)FS1425 | Mat1.2 | Cellule | vir |
|  | (INRA17-)FS1441 | Mat1.2 | Cellule | vir |
|  | (INRA17-)FS1477 | Mat1.1 | Cellule | vir |
|  | (INRA17-)FS1492 | Mat1.1 | Cellule | vir |
|  | (INRA18-)FS1721 | Mat1.2 | Apache | avr |
|  | (INRA18-)FS1738 | Mat1.2 | Apache | avr |
|  | (INRA18-)FS1754 | Mat1.1 | Apache | avr |
|  | (INRA18-)FS1775 | Mat1.1 | Apache | avr |
| 2019 | (INRA18-)FS2883 | Mat1.2 | Cellule | vir |
|  | (INRA18-)FS3098 | Mat1.2 | Cellule | vir |
|  | (INRA18-)FS2881 | Mat1.1 | Cellule | vir |
|  | (INRA18-)FS3250 | Mat1.1 | Cellule | vir |
|  | (INRA18-)FS2849 | Mat1.2 | Apache | avr |
|  | (INRA18-)FS2855 | Mat1.2 | Apache | avr |
|  | (INRA18-)FS2847 | Mat1.1 | Apache | avr |
|  | (INRA18-)FS2851 | Mat1.1 | Apache | avr |
| 2020 | (INRA19-)FS6176 | Mat1.2 | Cellule | vir |
|  | (INRA19-)FS6181 | Mat1.2 | Cellule | vir |
|  | (INRA19-)FS6141 | Mat1.1 | Cellule | vir |
|  | (INRA19-)FS6152 | Mat1.1 | Cellule | vir |
|  | (INRA19-)FS6503 | Mat1.2 | Apache | avr |
|  | (INRA19-)FS6522 | Mat1.2 | Apache | avr |
|  | (INRA19-)FS6191 | Mat1.1 | Apache | avr |
|  | (INRA19-)FS6196 | Mat1.1 | Apache | avr |

<sup>1</sup> determined by PCR amplification of the two mating type idiomorphs (Waalwijk *et al.*, 2002).
<sup>2</sup> based on the results of pathotyping on wheat seedlings, using cv. Cellule and Apache; ‘vir’ for virulent against *Stb16q* and ‘avr’ for avirulent against *Stb16q*.

**Table S2.** Intensity of asexual multiplication (disease severity) assessed for each type of co-inoculation and single inoculation in the three-year experiment. Disease severity is expressed as the mean percent sporulating area on the two uppermost leaves of wheat plants five weeks after inoculation. The impact of the type of co-inoculation on the intensity of asexual multiplication, by cultivar and year (with subsequent pooling of the three years) was assessed in Kruskal-Wallis tests with Bonferroni correction for pairwise comparisons. For single inoculations, the impact of strain status was assessed with Wilcoxon tests. Different letters indicate significant differences in means (*p* < 0.05) between statuses (columns) for a specific year × cultivar (rows) combination.

| Year | Cv. | Virulence status of the (co-) inoculated parental strains |  |  |  |  |
| --- | --- | --- | --- | --- | --- | --- |
| | | vir × vir | avr × vir | avr × avr | vir <sup>1</sup> | avr <sup>1</sup> |
| 2018 | Apache | 79.9 a | 77.1 a | 61.0 b | 59.8 $\alpha$ | 72.5 $\alpha$ |
| | Cellule | 30.5 a | 8.9 b | 0.1 c | 17.2 $\alpha$ | 0.1 $\beta$ |
| 2019 | Apache | 56.4 a | 58.4 a | 64.1 a | 33.8 $\alpha$ | 48.2 $\beta$ |
| | Cellule | 64.1 a | 24.1 b | 0.5 c | 17.4 $\alpha$ | 0.7 $\beta$ |
| 2020 | Apache | 35.2 a | 41.4 ab | 50.6 b | 48.3 $\alpha$ | 41.2 $\alpha$ |
| | Cellule | 38.7 a | 22.9 b | 0.1 c | 37.2 $\alpha$ | 0.0 $\beta$ |
| Means | Apache | 57.2 a | 59.0 a | 58.6 a | 47.3 $\alpha$ <sup>2</sup> | 54.0 $\alpha$ |
| 2018-2019-2020 | Cellule | 44.4 a | 18.6 b | 0.2 c | 23.9 $\alpha$ | 0.3 $\beta$ |

<sup>1</sup> single inoculation
<sup>2</sup> $p = 0.06$

**Table S3.** Genotyping data set (Excel file).

**Table S4.**
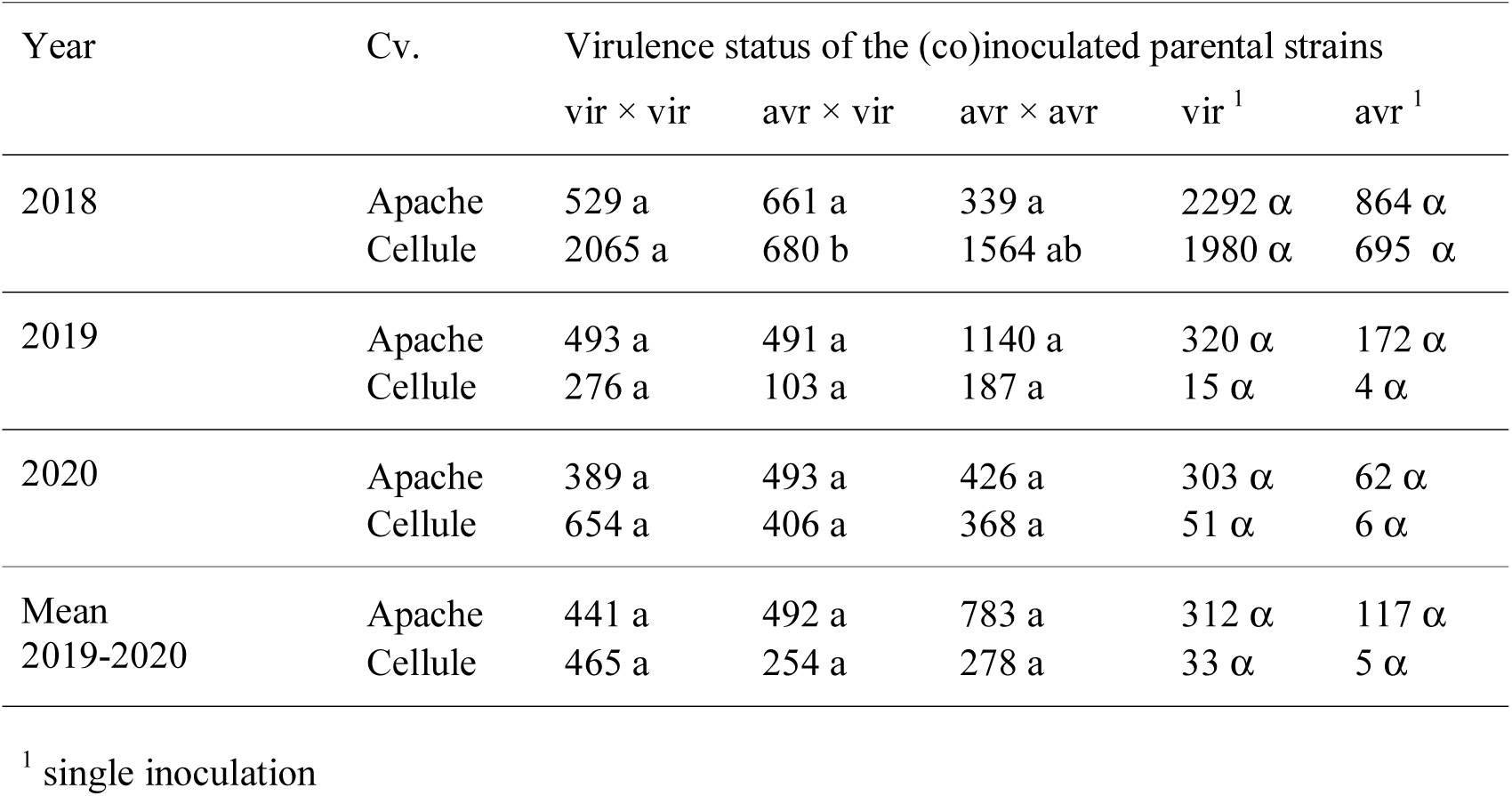
Intensity of sexual reproduction (ADI) assessed for each type of co-inoculation and single inoculation in the three-year experiment. ADI is expressed as the number of ascospores discharged per gramme of residues. The impact of the type of co-inoculation on the intensity of sexual reproduction for each cultivar and year (with subsequent pooling of the three years) was assessed in Kruskal-Wallis tests followed by Bonferroni correction for pairwise comparisons. For single inoculations, the impact of strain status was assessed with Wilcoxon tests. Different letters indicate significant differences in means (p < 0.05) between statuses (columns) for a specific year × cultivar (rows) combination.

| Year | Cv. | Virulence status of the (co)inoculated parental strains |  |  |  |  |
| --- | --- | --- | --- | --- | --- | --- |
| | | vir × vir | avr × vir | avr × avr | vir <sup>1</sup> | avr <sup>1</sup> |
| 2018 | Apache | 529 a | 661 a | 339 a | 2292 $\alpha$ | 864 $\alpha$ |
| | Cellule | 2065 a | 680 b | 1564 ab | 1980 $\alpha$ | 695 $\alpha$ |
| 2019 | Apache | 493 a | 491 a | 1140 a | 320 $\alpha$ | 172 $\alpha$ |
| | Cellule | 276 a | 103 a | 187 a | 15 $\alpha$ | 4 $\alpha$ |
| 2020 | Apache | 389 a | 493 a | 426 a | 303 $\alpha$ | 62 $\alpha$ |
| | Cellule | 654 a | 406 a | 368 a | 51 $\alpha$ | 6 $\alpha$ |
| Mean<br>2019-2020 | Apache | 441 a | 492 a | 783 a | 312 $\alpha$ | 117 $\alpha$ |
| | Cellule | 465 a | 254 a | 278 a | 33 $\alpha$ | 5 $\alpha$ |

<sup>1</sup> single inoculation

**Table S5.** Intensity of sexual reproduction (ADI) according to the proportion of the *Zymoseptoria tritici* virulent strain in the biparental inoculum suspension (‘avr × vir’). ADI is expressed as the number of ascospores discharged per gramme of residues of each cultivar (Apache, Cellule). Data were obtained in 2018, 2019 and 2020 for the FS1754 × FS1425, FS2883 × FS2847 and FS6176 × FS6191 co-inoculations, respectively. Differences in the mean intensity of sexual reproduction (pooling of the data for the three years) in a specific cultivar according to the proportion of the virulent parental strain were not significant (*p* > 0.05, Kruskal-Wallis test).

| Cv. | year | Proportion of the virulent parental strain |  |  |  |  |
| --- | --- | --- | --- | --- | --- | --- |
|  |  | 0.10 | 0.25 | 0.5 | 0.75 | 0.90 |
| Apache | 2018 | 638 | 1367 | 238 | 724 | 497 |
|  | 2019 | 302 | 59 | 665 | 318 | 1087 |
|  | 2020 | 1237 | 1340 | 398 | 273 | 1545 |
|  | mean | 726 | 922 | 434 | 438 | 1043 |
| Cellule | 2018 | 2043 | 883 | 375 | 48 | 1461 |
|  | 2019 | 82 | 3 | 167 | 111 | 0 |
|  | 2020 | 685 | 86 | 0 | 14 | 95 |
|  | mean | 937 | 324 | 181 | 57 | 518 |

| Cv. |  | Unbalanced ratio of parental strains |  |  |
| --- | --- | --- | --- | --- |
|  |  | 1/10 <sup>1</sup> | 1/4 <sup>2</sup> | 1/2 <sup>3</sup> |
| Apache | mean | 884 | 680 | 434 |
| Cellule | mean | 728 | 191 | 181 |

<sup>1</sup> 0.10-0.90 and 0.90-0.10
<sup>2</sup> 0.25-0.75 and 0.75-0.25
<sup>3</sup> 0.50-0.50

**Table S6.**
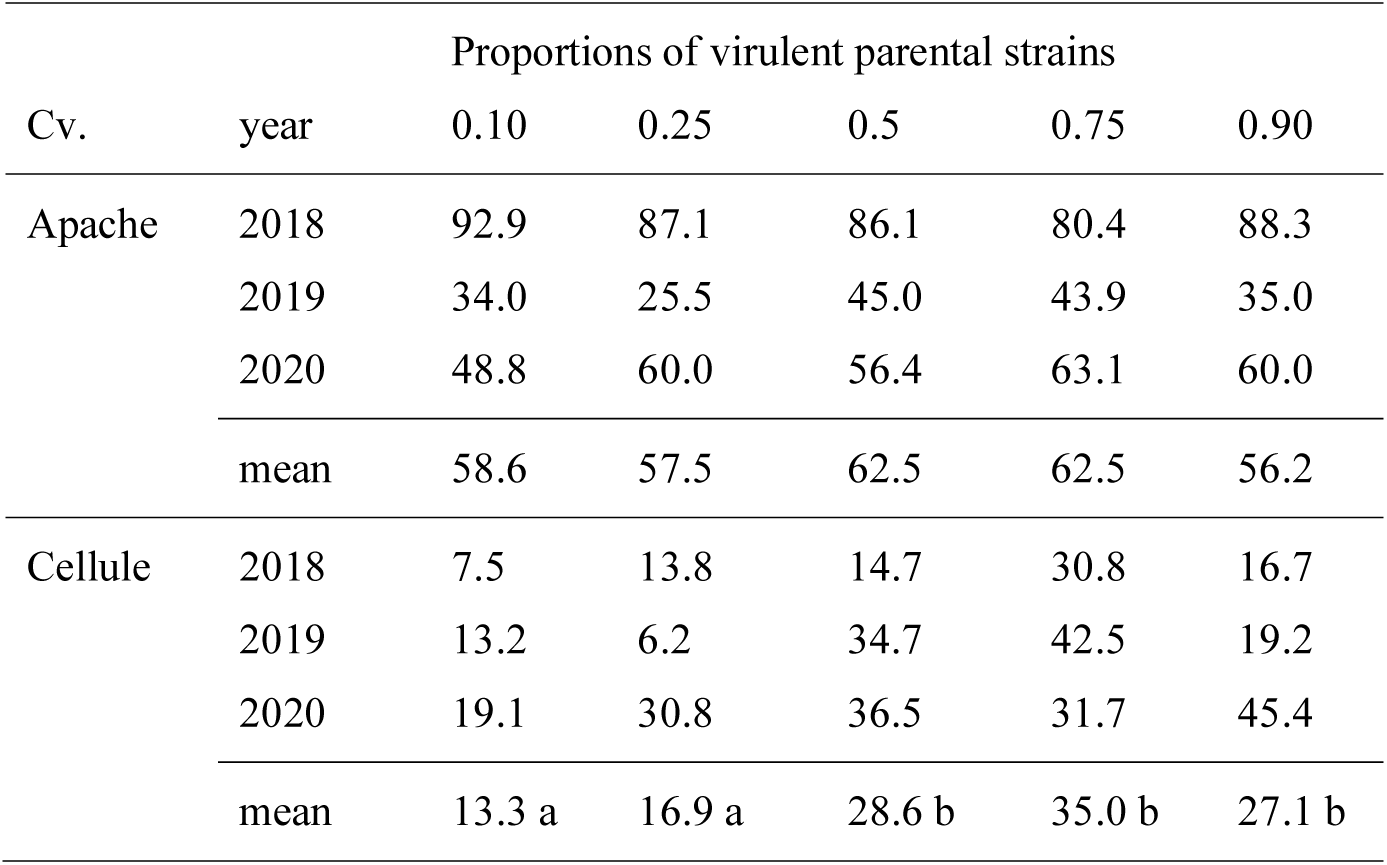
Intensity of asexual multiplication (disease severity) according to the proportion of the *Zymoseptoria tritici* virulent strain in the biparental suspension (‘avr × vir’). Disease severity is expressed as the mean percentage sporulating area on the two uppermost leaves of wheat plants five weeks after the inoculation of each cultivar (Apache, Cellule). Data were obtained in 2018, 2019 and 2020 for the FS1754 × FS1425, FS2883 × FS2847 and FS6176 × FS6191 co-inoculations, respectively. Different letters indicate significant differences in the mean intensity of asexual multiplication (pooling of the data for the three years) in a specific cultivar according to the proportion of the virulent parental strain (p < 0.01, Kruskal-Wallis test with Bonferroni correction for pairwise comparisons).

**Figure S1.**
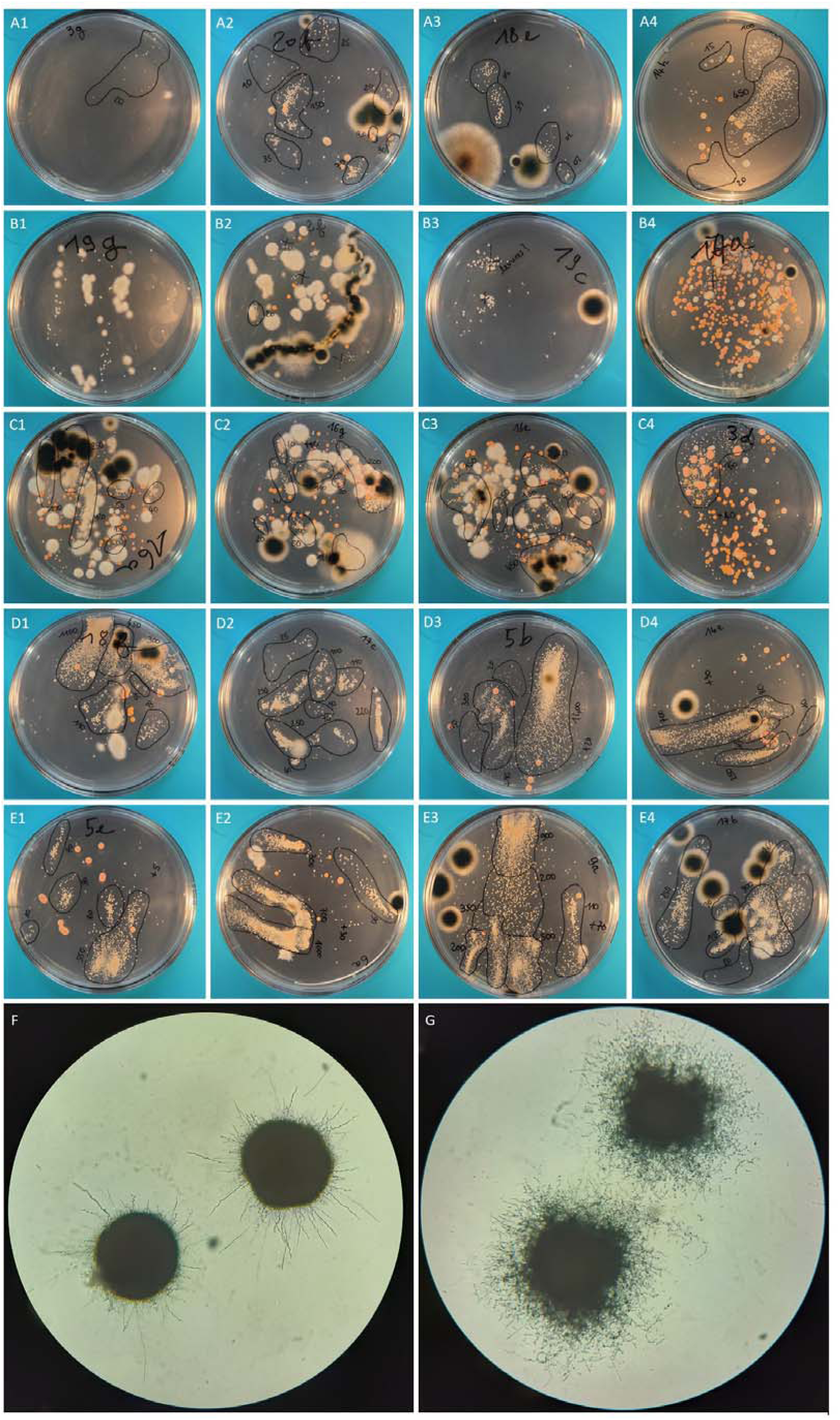
Clusters of *Zymoseptoria tritici* ascospore-derived colonies photographed seven days after ascospore ejection. **(A)** Little or no contamination; a few, distant, non-coalescent *Z. tritici* colonies of similar size, easy to identify and count. **(B)** Possible confusion between *Z. tritici* colonies (see **F**) and white bacterial or yeast colonies (see **G**). **(C)** Invasion of the plate by fast-growing fungi or bacteria covering all or part of the cluster of *Z. tritici* colonies. **(D)** Coalescence of *Z. tritici* colonies when cluster density is very high, precluding automated counting approaches based on image analysis. **(E)** High-density effect leading to different colony diameters, hindering automated area measurement approaches based on image analysis with software.

**Figure S2.**
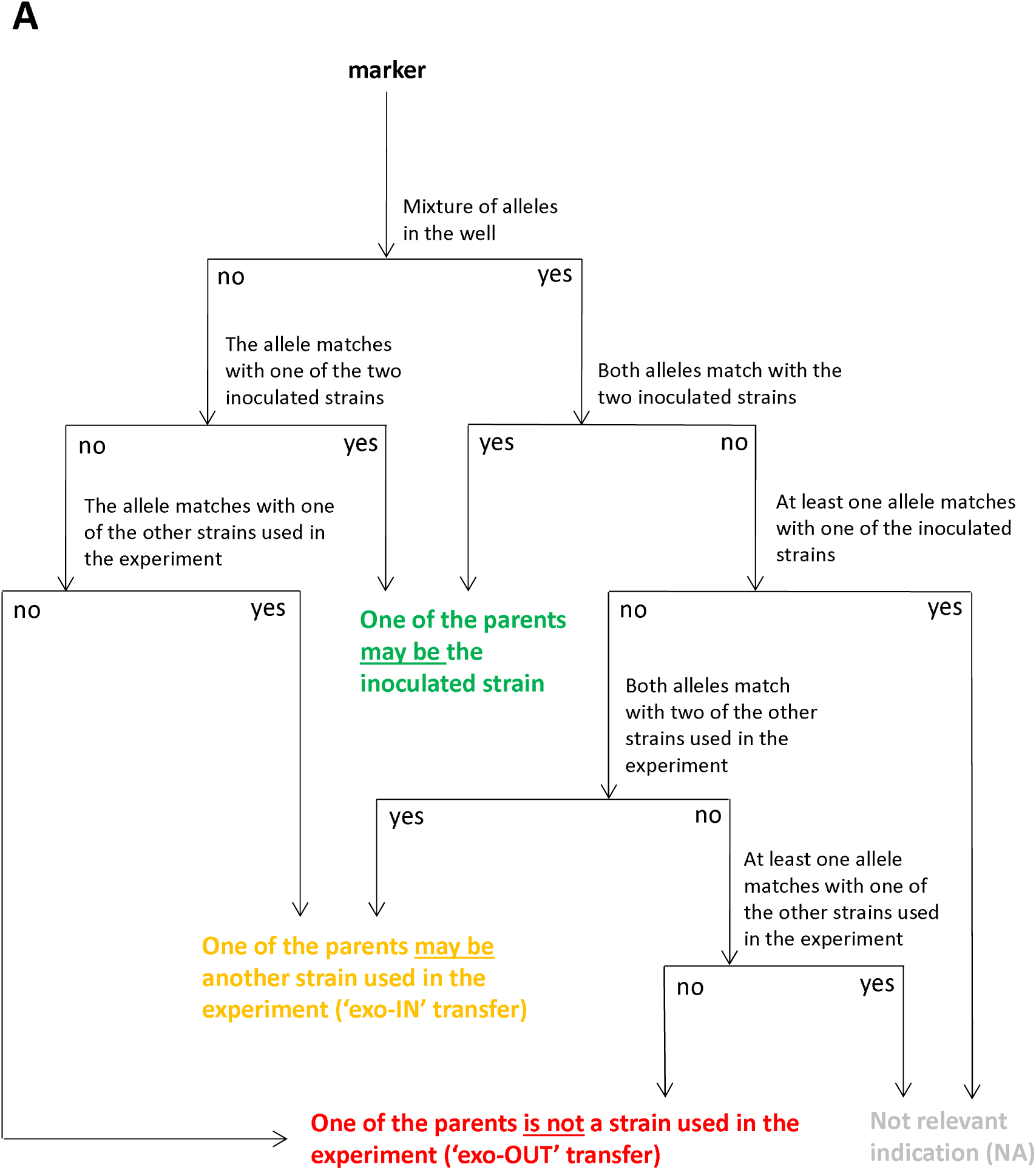
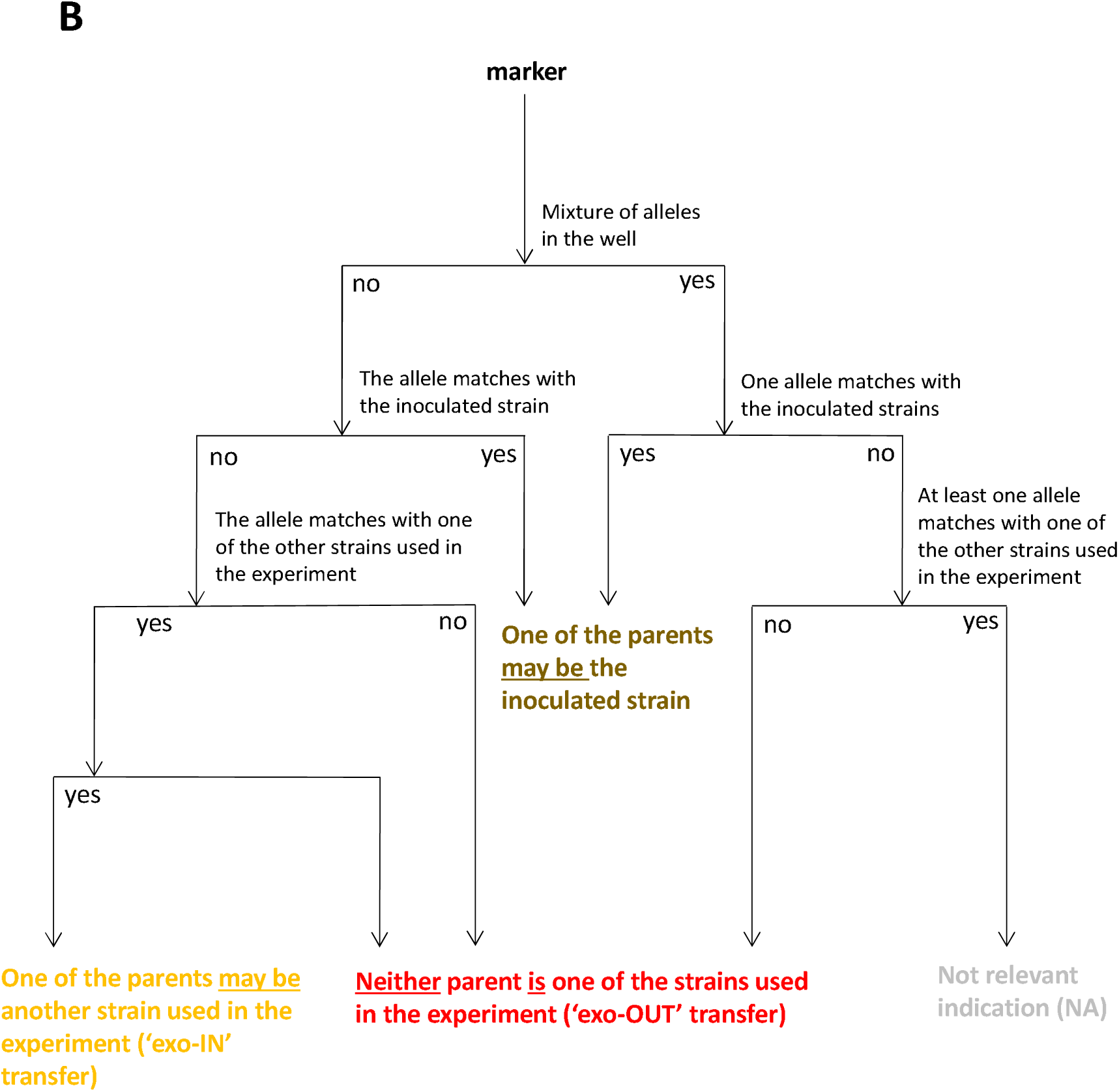
Decision trees used for investigation of the parenthood of *Zymoseptoria tritici* offspring strains based on the comparison of alleles for 12 SSR markers (m_i_) in the cases of (**A**) co-inoculation and (**B**) single inoculation. The color code is the one used for the genotyping data set in Table S3.

**Figure S3.**
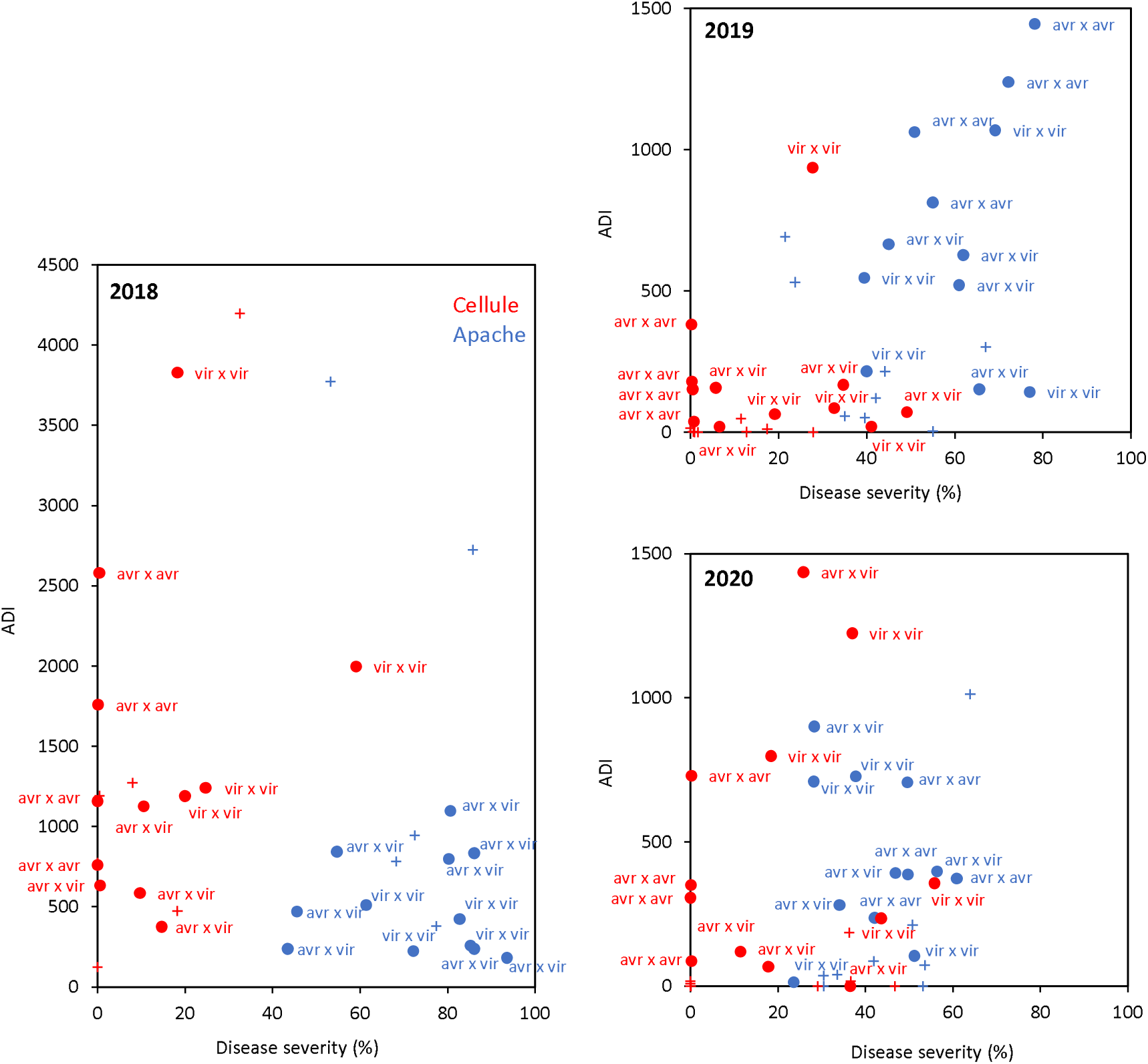
Intensity of *Zymoseptoria tritici* sexual reproduction (expressed as the number of ascospores discharged per gramme of residues of the same plant; ADI) and asexual multiplication (expressed as disease severity), by cultivar (Apache, Cellule) and type of inoculation during the three-year crossing experiment. The number of isolates used for inoculation (+ for single inoculations, • for co-inoculations), their virulence status with respect to *Stb16q* (‘v’ for a virulent strain, ‘a’ for an avirulent strain) and the wheat cultivar (red for Cellule, blue for Apache) are indicated for each point, with the data averaged for nine stems (three plants). There was no correlation between the intensity of sexual reproduction and the intensity of asexual multiplication (R² = 0.38).

